# Identification of Physiological Response Functions to Correct for Fluctuations in Resting-State fMRI related to Heart Rate and Respiration

**DOI:** 10.1101/512855

**Authors:** Michalis Kassinopoulos, Georgios D. Mitsis

**Author notes:** Corresponding author: Department of Bioengineering, McGill University, 350 McConnell Engineering Building, 3480 University Street, Montreal, QC, Canada H3A 0E9, E-mail address (G.D. Mitsis).

## Abstract

Functional magnetic resonance imaging (fMRI) is widely viewed as the gold standard for studying brain function due to its high spatial resolution and non-invasive nature. However, it is well established that changes in breathing patterns and heart rate strongly influence the blood oxygen-level dependent (BOLD) fMRI signal and this, in turn, can have considerable effects on fMRI studies, particularly resting-state studies. The dynamic effects of physiological processes are often quantified by using convolution models along with simultaneously recorded physiological data. In this context, physiological response function (*`PRF`*) curves (cardiac and respiratory response functions), which are convolved with the corresponding physiological fluctuations, are commonly employed. While it has often been suggested that the *`PRF`* curves may be region- or subject- specific, it is still an open question whether this is the case. In the present study, we propose a novel framework for the robust estimation of *`PRF`* curves and use this framework to rigorously examine the implications of using population-, subject-, session- and scan-specific *`PRF`* curves. The proposed framework was tested on resting-state fMRI and physiological data from the Human Connectome Project. Our results suggest that *`PRF`* curves vary significantly across subjects and, to a lesser extent, across sessions from the same subject. These differences can be partly attributed to physiological variables such as the mean and variance of the heart rate during the scan. The proposed methodological framework can be used to obtain robust scan-specific *`PRF`* curves from data records with duration longer than 5 minutes, exhibiting significantly improved performance compared to previously defined canonical cardiac and respiration response functions. Besides removing physiological confounds from the BOLD signal, accurate modeling of subject- (or session-/scan-) specific *`PRF`* curves is of importance in studies that involve populations with altered vascular responses, such as aging subjects.

**Highlights:**

- Physiological response functions (*`PRF`*) vary considerably across subjects/sessions
- Scan-specific *`PRF`* curves can be obtained from data records longer than 5 minutes
- The shape of the cardiac response function is linked to the mean heart rate (HR)
- Brain regions affected by HR and breathing patterns exhibit substantial overlap
- HR and breathing patterns affect distinct regions as compared to cardiac pulsatility

## 1. Introduction

Over the last few decades, advances in neuroimaging methods have significantly facilitated the study of brain function. One of the most popular methods for measuring brain activity is functional magnetic resonance imaging (fMRI), due to its high spatial resolution and non-invasive nature. fMRI, in principle, allows the mapping of brain function by measuring the hemodynamic response that accompanies neuronal activity in the brain. The onset of neuronal activity leads to physiological changes in the cerebrovascular system, including changes in cerebral blood flow (CBF), cerebral blood volume (CBV) per unit of brain tissue, as well as oxyhemoglobin and deoxyhemoglobin concentrations. The majority of fMRI studies is based on the blood oxygen-level dependent (BOLD) contrast mechanism, which primarily corresponds to the concentration of deoxyhemoglobin, and, thus, reflects a complex interplay of the aforementioned physiological processes (Ogawa et al., 1990).

Intriguingly, low-frequency (< 0.15 Hz) fluctuations of the BOLD fMRI signal in the resting brain have consistently revealed significant correlations between distinct brain regions giving rise to a number of functional networks, termed resting-state networks (RSNs) (Biswal et al., 1995; Smith et al., 2009). Furthermore, several studies have reported alterations of RSNs in a range of cerebrovascular and mental disorders, demonstrating their potential use as biomarkers (Demirtaş et al., 2016; Leonardi et al., 2013; Sheline et al., 2010; Woodward and Cascio, 2015). Therefore, while fMRI studies were initially focused on studying the function of individual brain regions in response to specific tasks, during the last two decades or so there has been a shift towards understanding the correlation between distinct brain regions during rest, referred to as resting-state functional connectivity (rs-FC; van den Heuvel and Hulshoff Pol, 2010).

However, the interpretation of rs-FC studies is often questioned, partly because of the challenge of disentangling the neuronal component of the BOLD signal, which is typically of interest, from measurement and physiological confounds (Bright and Murphy, 2015; Murphy et al., 2009). These confounds may be related to scanner hardware drifts and instabilities, head motion as well as spontaneous physiological fluctuations, including breathing, cardiac activity and arterial CO_2_ (Caballero-Gaudes and Reynolds, 2017; Liang et al., 2015; Murphy et al., 2013; Wise et al., 2004). For instance, Fluctuations in the BOLD signal arise from cardiac pulsation, which pushes the brainstem into the surrounding brain tissue, causing deformation and cerebrospinal fluid movement (Dagli et al., 1999) while breathing-induced fluctuations result partly from breathing-related bulk movement of the head (Hu et al., 1995). Furthermore, variations in the air volume inside the lungs as well as chest movement due to breathing lead to changes in the static magnetic field which, in turn, cause an image shift in the phase encoding direction, consequently distorting the fMRI volumes (Bollmann et al., 2017; Raj et al., 2001). Importantly, variations in the rate or depth of breathing have an impact on the arterial tension of CO_2_, which is a potent vasodilator and can therefore induce changes in CBF (Birn et al., 2006; Wise et al., 2004). In turn, global CBF changes cause low-frequency (~0.1 Hz) fluctuations in the BOLD signal, which may be misinterpreted as neuronal activity (Birn et al., 2008, 2006). In addition, it has been shown that fluctuations in the BOLD signal are caused by spontaneous fluctuations in heart rate (HR; Napadow et al., 2008; Shmueli et al., 2007). These physiological-related fluctuations can have considerable impact on the resulting rs-FC patterns, including dynamic rs-FC patterns, as they tend to inflate the correlation between areas affected by physiological noise (Birn et al., 2008; Nikolaou et al., 2016). Therefore, several physiological noise correction techniques have been developed to remove the effects of physiological factors from fMRI data.

One of the most widely used methods for fMRI physiological noise correction is RETROICOR, proposed by Glover et al. (2000). According to this method, the pulsatility of blood flow and breathing motion are considered to distort the BOLD signal inducing an artifact that is time-locked to the cardiac and respiratory phases. Therefore, the associated physiological regressors are estimated as a linear combination of sinusoidal signals coinciding with the cardiac and breathing cycles using concurrent cardiac and respiratory measurements and subsequently regressed out. RETROICOR can effectively remove the high-frequency cardiac (~1 Hz) and breathing (~0.3 Hz) artifacts, despite the aliasing that takes place in a typical fMRI acquisition with a relatively low sampling rate (e.g. TR=3 s; Jones et al., 2008).

Cardiac and respiratory recordings can be also used for reducing low-frequency BOLD fluctuations associated with changes in HR and breathing patterns using physiological response function (*`PRF`*) models, such as the ones proposed by Chang et al. (2009) and Birn et al. (2006, 2008). In the first model, HR values, extracted from cardiac measurements, are convolved with the so-called cardiac response function (*`CRF`*). According to the second model, respiration volume per time (RVT), which is a measure proportional to the breathing rate (BR) and depth at each time point, is initially estimated based on measurements from a pneumatic belt. Subsequently, RVT is convolved with the respiration response function (*RRF*) to estimate BOLD fluctuations due to changes in the breathing pattern. Both models are implemented in major fMRI preprocessing toolboxes such as the physiological noise modelling (PNM) toolbox of FSL (Jenkinson et al., 2012) and the PhysIO SPM toolbox (Kasper et al., 2017). Nevertheless, their use has been somewhat limited, partly due to that they do not account for between-subject *`PRF`* variability. Interestingly, in this context, Falahpour et al. (2013) proposed an alternate approach for constructing subject-specific *`PRF`* curves based on the global signal (GS), which is defined as the mean BOLD signal across all voxels in the brain, from each scan. Physiological regressors constructed in this way can account for a considerably larger fraction of variance in fMRI time-series compared to the standard *`PRF`* curves. However, when individual *`PRF`* curves were used in a cross-validation analysis, the results suggested that the improvement in the explained variance may be due to overfitting (Falahpour et al., 2013).

Several data-driven approaches have been also proposed for preprocessing BOLD fMRI data. For example, in global signal regression (GSR), the GS is subtracted from the data through linear regression, implicitly assuming that processes that globally affect the fMRI BOLD signal are mostly uncorrelated to neural activity (Fox et al., 2005; Greicius et al., 2003; Qing et al., 2015). Power et al. (2017) have recently shown that statistical maps demonstrating the correlation between each voxel time-series and the GS resemble statistical maps reported in Wise et al. (2004) with areas affected by fluctuations in arterial CO_2_, while Liu et al. (2017) have identified a frequency range (0.02-0.04 Hz) in the GS that acts as a proxy for spontaneous changes in CO_2_. While these findings may be in support of GSR in order to correct for breathing-related changes in CO_2_, the validity of GSR is still under debate, as there is some evidence that the GS has a neuronal component as well (Liu et al., 2017; Murphy and Fox, 2017). Furthermore, the use of independent component analysis (ICA) or principal component analysis (PCA) to identify physiological or “noisy” components (based on their temporal, spatial and spectral features) and subsequently remove them before reconstructing the “noise-free” fMRI data, has been proposed (Churchill and Strother, 2013; Kay et al., 2013; Pruim et al., 2015; Salimi-Khorshidi et al., 2014). Even though the performance of data-driven approaches depends on subjective choices (e.g. the number of components to be removed when performing aCompCor (Behzadi et al., 2007) or the selection of the noisy components during the training phase when using FIX), these approaches have the advantage that they do not require collection of physiological data, which is often omitted in fMRI studies. FIX (“FMRIB’s ICA-based X-noisefier”) is a data-driven technique, proposed by Salimi-Khorshidi et al. (2014), which implements semi-automatic procedure for denoising fMRI via classification of ICA components. Due to its performance with respect to automatic and manual classification of “noisy” components, FIX has been used in the default resting-state fMRI preprocessing pipeline for generating HCP connectomes (Salimi-Khorshidi et al., 2014). However, recent studies have demonstrated that global fluctuations captured in the GS are still prominent after FIX-denoising (Burgess et al., 2016; Power et al., 2017). Additional studies have established a strong association of the GS with slow-frequency fluctuations of breathing and heart rate (Chang and Glover, 2009a; Falahpour et al., 2013). Overall, these studies suggest that FIX, and in general PCA/ICA-based noise correction techniques, may not sufficiently correct for these noise sources.

In the present paper, we propose a novel methodological framework for extracting subject- and scan-specific *`PRF`* curves using the GS. We propose a double gamma structure for the *`PRF`* curves and a combination of optimization techniques (genetic algorithms and interior-point optimization) for parameter estimation. In contrast to previous approaches (Birn et al., 2008; Chang et al., 2009; Falahpour et al., 2013), the convolution of physiological variables (HR, breathing pattern) with the *`PRF`* curves is done in a pseudo-continuous time-domain at a 10 Hz sampling frequency to avoid smoothing out the effect of high-frequency physiological fluctuations of HR. In addition to between-subject variability, we rigorously investigate the between-session variability of the *`PRF`* curves, as well as their variability across voxels in the brain. To this end, as well as to evaluate the performance of the proposed methodology, we use resting-state fMRI data from the Human Connectome Project (HCP; Van Essen et al., 2013) collected during 4 different scans on 2 different days. The noise correction techniques discussed here are also of importance for task-based studies, such as those involving motor and pain protocols, as fluctuations in cardiac activity and breathing patterns may be time-locked to the task, and, hence, bias the results (Glasser et al., 2018). The codes for the *`PRF`* models proposed here are publicly available and can be found on https://github.com/mkassinopoulos/PRF_estimation, while the group-level statistical maps derived from this dataset with the brain regions affected by cardiac and breathing activity are available on NeuroVault (Gorgolewski et al., 2015) for detailed inspection (https://neurovault.org/collections/5654/).

## 2. Methodology

### 2.1 Human Connectome Project (HCP) Dataset

We used resting-state scans from the HCP S1200 release (Glasser et al., 2016; Van Essen et al., 2013). The HCP dataset includes, among others, resting-state (eyes-open and fixation on a cross-hair) data from healthy young (age range: 22-35 years) individuals acquired on two different days. On each day, two 15-minute scans were collected. We refer to the two scans collected on days 1 and 2 as R1a/R1b and R2a/R2b respectively. fMRI acquisition was performed with a multiband factor of 8, spatial resolution of 2 mm isotropic voxels, and a repetition time TR of 0.72 s (Glasser et al., 2013).

The minimal preprocessing pipeline for the resting-state HCP dataset is described in (Glasser et al., 2013). In brief, the pipeline includes gradient-nonlinearity-induced distortion correction, motion correction, EPI image distortion correction, non-linear registration to MNI space and mild high-pass (2000 s) temporal filtering. The motion parameters are included in the database for further correction of motion artifacts. The HCP has adopted FIX for removing structured temporal noise related to motion, non-neuronal physiology, scanner artefacts and other nuisance sources (Salimi-Khorshidi et al., 2014). FIX-denoised data are available in the HCP database as well.

In the present work, we examined minimally-preprocessed and FIX-denoised data from 41 subjects (Supplementary Table 1), which included good quality physiological signals (cardiac and breathing waveforms) in all four scans, as assessed by visual inspection. The cardiac and respiratory^1^ signals were collected with a photoplethysmograph and respiratory belt respectively.

### 2.2 Preprocessing

Unless stated otherwise, the preprocessing and analysis described below were performed in Matlab (R2017b; Mathworks, Natick MA).

#### 2.2.1 Preprocessing of physiological recordings

The cardiac signal (i.e. photoplethysmogram) was initially band-pass filtered with a 2^nd^ order Butterworth filter between 0.3 and 10 Hz. The minimum peak distance specified for peak detection varied between 0.5 and 0.9 seconds depending on the subject’s average HR. The HR signal was computed in beats-per-minute (bpm) by multiplying the inverse of the time differences between pairs of adjacent peaks with 60, and evenly resampled at 10 Hz.

In the case of scans with abrupt changes in HR, if these changes were found by visual inspection to be due to noisy cardiac signal measurements, the HR signal was corrected for outliers using Matlab. Specifically, outliers in the HR signal were defined as the time points that deviated more than (approximately) 7 median absolute deviations (MAD) from the moving median value within a time window of 30 seconds. The MAD threshold varied across scans and was chosen empirically based on the extent of noise in the cardiac signal of each scan and the extracted HR signal. Outliers were replaced using linear interpolation (for examples of HR signals with abrupt changes and how they were treated, please see Supplementary Figs. 1 and 2).

The respiratory signal was detrended linearly and corrected for outliers using the moving median method described earlier. The duration of the moving time window and the MAD threshold were chosen separately for each scan based on visual inspection. Subsequently, the respiratory signal was low-pass filtered at 5 Hz with a 2^nd^ order Butterworth filter and z-scored. The peak detection, used later for the extraction of RVT, was done with a minimum peak distance of 2 s and minimum peak height of 0.2. The figures of the physiological recordings, before and after the preprocessing, as well as the HR and breathing-related features can be viewed through our figshare repository (https://doi.org/10.6084/m9.figshare.c.4585223.v4; Kassinopoulos and Mitsis, 2019).

#### 2.2.2 Preprocessing of fMRI data

The effect of HR and respiratory variations on the fMRI BOLD signal is considered to last about half a minute (Chang et al., 2009). Therefore, the first 40 image volumes were disregarded, while the corresponding physiological data were retained. The fMRI time-series were first spatially smoothed with a Gaussian filter of 3 mm full width at half maximum (FWHM) and then linearly detrended. Subsequently, the following nuisance variables were regressed out through linear regression: the demeaned and linearly detrended motion parameters and their derivatives, and 3^rd^ order RETROICOR regressors for the cardiac and respiratory artifacts (Glover et al., 2000) using the physiological recordings at the original sampling rate of 400 Hz.

### 2.3 Physiological response functions

We employed linear dynamic models for extracting physiological regressors that were subsequently included in the general linear model as regressors to model the effect of the corresponding physiological variable on the BOLD signal. The physiological regressors were obtained as the convolution between the physiological variables and the corresponding *`PRF`*.

#### 2.3.1 Standard cardiac response function (*CRF*_*stand*_; Chang et al., 2009)

In the present study, the model proposed in Chang et al. (2009) was considered as the standard method for removing the effect of HR fluctuations. According to this method, the HR signal is smoothed with a 6 *s* moving average filter before being convolved with the standard *`CRF`* (*`CRF`*_*stand*_) defined as:

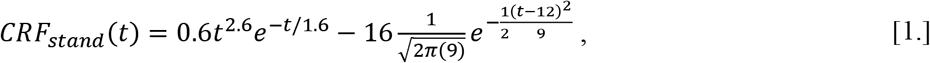

to construct the physiological regressor *X*_*HR*_ related to HR fluctuations. Finally, *X*_*HR*_ was downsampled to the fMRI sampling rate.

#### 2.3.2 Standard respiration response function (RRF_stand_; Birn et al., 2008, 2006)

We used the method described in (Birn et al., 2008) as the standard method for removing the effect of changes in the breathing pattern. Briefly, the maximum and minimum peaks of each breath were initially identified on the respiratory signal and linearly interpolated at 10 Hz. Subsequently, the breathing depth was defined as the difference between the interpolated time-series of maximum and minimum peaks. The BR was calculated as the time difference between successive maximum peaks, expressed in respirations-per-minute (rpm), and interpolated at 10 Hz. Subsequently, the respiration volume per time (RVT) was calculated as the product between breathing depth and rate. Then, the RVT time-series was convolved with the standard *RRF* (*RRF*__*stand*__) defined as:

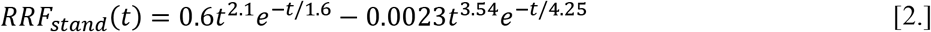

which yielded the physiological regressor *X*_*RVT*_ related to changes in the breathing pattern. Finally, *X*_*RVT*_ was downsampled to the fMRI sampling rate.

#### 2.3.3 Proposed physiological response functions (*PRF*)

Here, we used the instantaneous HR without any smoothing, whereas the respiratory signal was first filtered with a moving average window of 1.5 s and the square of its derivative was subsequently calculated. The respiratory signal was smoothed before calculating the derivative to avoid large, physiologically implausible spikes in the extracted regressor. The signal extracted after this process, termed respiratory flow (RF), reflects the absolute flow of the inhalation and exhalation of the subject at each time point. While RF carries similar information to RVT, it is expected to be more robust as it does not depend on accurate peak detection. To reduce the time of subsequent analysis, RF was downsampled from 400 Hz to 10 Hz. Therefore, the corresponding physiological regressors were defined as follows:

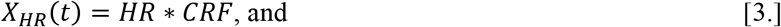

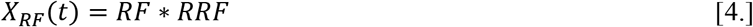

where *`CRF`* and *RRF* are the proposed cardiac and respiration response functions, respectively. The basic structure of the two proposed *`PRF`* curves was selected as the double gamma function that is also used for *RRF*__*stand*__ and the canonical hemodynamic response function (HRF) in the SPM software package (http://www.fil.ion.ucl.ac.uk/spm/). The gamma function is defined as:

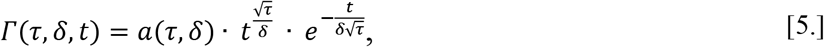

The parameters τ and δ indicate the (approximate) time of peak and dispersion of the function, and the parameter α is a scaling factor which normalizes the peak value of the gamma function to 1. The *`PRF`* curves are defined as follows:

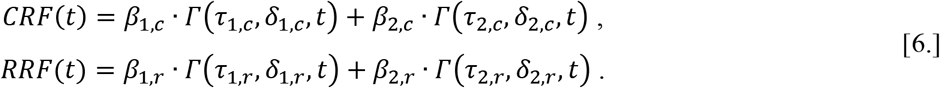

Below, we collectively refer to the eight parameters of the gamma functions (τ_1,*c*_, δ_1,*c*_, τ_2,*c*_, δ_2,*c*_, τ_1,*r*_, δ_1,*r*_, τ_2,*r*_, δ_2,*r*_) and the four scaling parameters (*β*_1,*c*_, *β*_2,*c*_, *β*_1,*r*_, *β*_2,*r*_) as ***G*** and ***B***, respectively. Note that, since the *`PRF`* curves have arbitrary units, they can be expressed as follows:

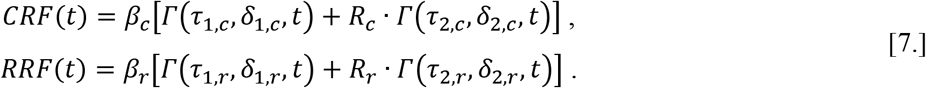

where the parameters *R*_*c*_ and *R*_*r*_ correspond to the ratios *β*_2,*c*_/*β*_1,*c*_ and *β*_2,*r*_/*β*_1,*r*_, respectively, and the two scaling parameters *β*_*c*_ and *β*_*r*_ reflect the amount of variance explained by the corresponding physiological variables on the BOLD signal. Finally, the extracted physiological regressors were downsampled to the fMRI acquisition rate.

### 2.4 Comparison of different physiological models

Previous studies have suggested that subject-specific *`PRF`* curves may be more appropriate for constructing physiological regressors (Birn et al., 2008; Chang et al., 2009; Falahpour et al., 2013). Here, we rigorously examined this hypothesis by considering several different cases for estimating the *`PRF`* curve parameters and assessing the resulting performance. For each subject, we used four different 15-minute resting-state scans collected on two different sessions (days): R1a/R1b (day one) and R2a/R2b (day two). This allowed us to examine the variability of the *`PRF`* curves between subjects, as well as between scans and sessions of the same subject. Initially (Section 2.4.1), we considered three main models (two variants of population-specific models and a scan-specific model) to examine the variability in the shape of the *`PRF`* curves across scans for models with different degree of flexibility. Subsequently (Section 2.4.2), we assessed the performance of several variants of *`PRF`* models with respect to the explained variance to examine whether the use of subject-, session- or scan- specific *`PRF`* is justifiable. In both cases, the GS from each scan was used to define the *`PRF`* curves and assess model performance. To extend these results (Section 2.4.3), we compared the performance of a subset of the models considered in 2.4.2 as well as the performance of a voxel-specific model in individual voxels.

#### 2.4.1 Variability in the shape of the *PRF* curves across scans

Here, we aimed to examine the variability in the shape of *`PRF`* curves across scans for models based on different degree of flexibility. In addition, we aimed to understand the relation of the variability in shape to physiological variables such as the mean HR. To this end, we examined three variants of the proposed *`PRF`* curves, termed *`PRF`*_*pop*_, 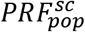 and *`PRF`*_*sc*_ that were used to explain fluctuations on the GS of each scan. The *`PRF`*_*pop*_ population-specific model is based on Eq. 7 and is the least flexible model, as it assumes that ***G*** (i.e. τ_1,*c*_, δ_1,*c*_, τ_2,*c*_, δ_2,*c*_, τ_1,*r*_, δ_1,*r*_, τ_2,*r*_, δ_2,*r*_) and ***R*** (i.e. *R*_*c*_, *R*_*r*_) are the same for all subjects. In this model, only ***B*** (i.e. *β*_*c*_, *β*_*r*_), which determines the amount of variance explained by HR and breathing pattern on the GS for each scan, was allowed to vary across scans. The 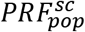 model is also a population-specific model that allows variability in the shape of the *`PRF`* curves between scans. Specifically, it is based on Eq. 6 and it assumes that ***G*** (τ_1,*c*_, δ_1,*c*_, τ_2,*c*_, δ_2,*c*_, τ_1,*r*_, δ_1,*r*_, τ_2,*r*_, δ_2,*r*_) is the same for all subjects while ***B*** (i.e. *β*_1,*c*_, *β*_2,*c*_, *β*_1,*r*_, *β*_2,*r*_) was allowed to vary across scans. However, in this case, ***B*** determines both the amount of variance explained by the physiological variables on the GS as well as the shape of the *`PRF`* curves. Finally, *`PRF`*_*sc*_ is a scan-specific model also based on Eq. 6, in which all the parameters were allowed to vary across scans. The *`PRF`* parameters were estimated with the non-linear optimization techniques described in Section 2.4.1.1.

In the case of the scan-specific model *`PRF`*_*sc*_, apart from comparing the curves with their standard counterparts and population-specific curves *`PRF`*_*pop*_, we also examined whether the physiological variables for each scan can explain the between-scan variability in the shape of the curves as well as the performance with respect to the explained variance on the GS. Specifically, we examined whether the mean and variance of HR and BR/RF were correlated with the time of positive and negative peaks for *`CRF`*_*sc*_ and *RRF*_*sc*_, respectively. In addition, we examined whether the mean and variance of HR and BR/RF were correlated to the correlation coefficient values between the corresponding physiological regressors (Eqs. 3-4) and the GS. Overall, we examined the relationship of 18 pairs of variables consisting of 6 explanatory variables, the mean and variance of HR, BR and RF, and 6 dependent variables, the time of positive and negative peak of the *`CRF`*_*sc*_/*RRF*_*sc*_ curves as well as the correlation coefficient between the corresponding physiological regressors and the GS. Pairs of variables related to both HR and breathing pattern (e.g. mean HR and time of positive peak for *RRF*_*sc*_) were not considered. Regarding statistical testing, we used an alpha level of .05 adjusted for multiple comparisons (N=18) with Bonferroni correction.

##### 2.4.1.1 PRF parameter estimation

The parameters of the population-specific model *`PRF`*_*pop*_ were estimated as follows (for a pseudocode of the algorithm, please see Supplementary Table 2): 1. for a set of given parameters, the two *`PRF`*_*pop*_ curves were constructed. Subsequently, for each scan, 2. the HR and RF signals (sampled at 10 Hz) were convolved with *`CRF`*_*pop*_ and *RRF*_*pop*_ respectively to extract the corresponding physiological regressors and then downsampled to match the fMRI acquisition rate. 3. Estimation of the general linear model (GLM) was performed, whereby the GS was the dependent variable and the two physiological regressors were the two explanatory variables. 4. the Pearson correlation coefficient between the GS and the model prediction was calculated. 5. after performing steps 1-4 for all scans, the correlation value was averaged across all scans and returned by the algorithm and 6. the parameter values that maximized this correlation value were estimated using numerical optimization techniques as described later.

In the case of 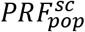, steps 2 and 3 were implemented as follows: In step 2, the two gamma functions for each 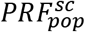 curve were convolved separately with HR and RF. As a result, in step 3, two physiological regressors related to HR and two regressors related to RF were included in the GLM as explanatory variables (for a pseudocode of the algorithms for the *`PRF`*_*pop*_ and 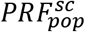 models, please see Supplementary Table 2). Finally, the procedure described above was followed for the estimation of the parameters in the scan-specific model *`PRF`*_*sc*_, with the only difference being that step 5 was omitted, as the *`PRF`*_*sc*_ parameters were estimated separately for each scan.

To obtain the optimal parameter values for the *`PRF`*_*pop*_ model, a genetic algorithm (GA) implemented in Matlab R2017b’s Global Optimization Toolbox was initially applied. GAs are a family of popular heuristic optimization techniques that search the parameter space for the optimal solution of a problem in a manner inspired by Darwin’s principle of natural selection (Holland, 1975). GAs have generally higher demands in CPU time compared to gradient-based algorithms, but they are capable of providing potentially global optimal solutions for complex functions (Patel and Padhiyar, 2015). The parameter vectors ***τ*** (τ_1,*c*_, τ_2,*c*_ τ_1,*r*_, τ_2,*r*_) and ***δ*** (δ_1,*c*_, δ_2,*c*_, δ_1,*r*_, δ_2,*r*_) were bounded between 0-20 seconds and 0-3 seconds, respectively. A stopping criterion of 100 generations was set, as it was found to be adequate for convergence. The solution of the GA was subsequently used as the initial point for the interior-point gradient-based algorithm, also implemented in Matlab R2017b (Optimization Toolbox), with a stopping criterion of 100 maximum iterations, to refine the solution. To accelerate the estimation procedure for the remaining models, rather than using the GA for initialization of the parameters, the obtained *`PRF`*_*pop*_ parameter values (or curves) were used as the initial point for all models and the interior-point algorithm was employed with a stopping criterion of 100 maximum iterations to refine the solution. Moreover, since the parameter estimation for the scan-specific model was performed using a smaller amount of data, making this model more prone to overfitting, the upper and lower boundaries for ***τ*** and ***δ*** were restricted within a 6 sec range centered around the population-specific *`PRF`*_*pop*_ model parameter values. In case of negative lower boundaries, these values were set to zero.

#### 2.4.3 Comparison of population-, subject-, session- and scan-specific *PRF* curves

This section aimed to assess the performance of 13 different models (Table 1) with respect to the explained variance using two-level cross-validation (CV) to examine whether *`PRF`* curves significantly vary between subjects as well as between sessions or scans within-subject. For each model, the *`PRF`* parameters were estimated from one segment of data (training set at the first-level of CV; 3_rd_ column of Table 1) and model performance was assessed in a separate segment of data (validation set at the first-level of CV; 4^th^ column of Table 1) as described in sections 2.4.2.1 and 2.4.2.2, respectively.

**Table 1.**
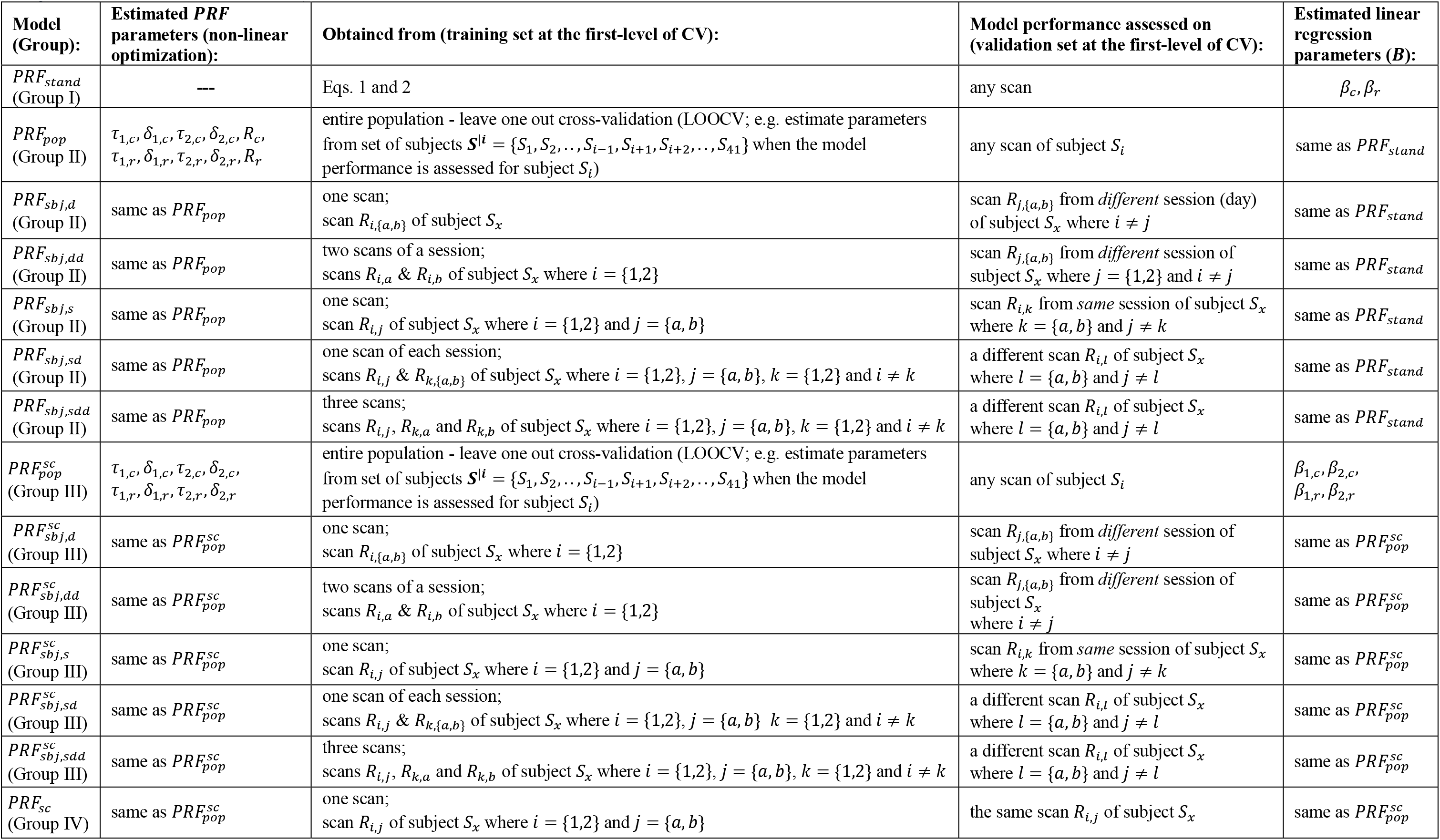
Assessment of the performance of population-, subject-, session- and scan-specific `PRF` models using two-level cross-validation (CV; *stand*: standard, *ppl*: population, *sbj*: subject, *sc*: scan, *d*: different, *s*: same)

##### 2.4.2.1 PRF parameter estimation at the first-level of cross-validation

The *`PRF`* models listed in Table 1 are, to some extent, sorted by the least to the most flexible model. The first six models (*`PRF`*_*pop*_ to *`PRF`*_*sbj,sdd*_, group of models II) are based on Eq. 7. In this case, ***R*** (*R*_*c*_, *R*_*r*_), along with ***G*** (τ_1,*c*_, δ_1,*c*_, τ_2,*c*_, δ_2,*c*_, τ_1,*r*_, δ_1,*r*_, τ_2,*r*_, δ_2,*r*_), define the shape of the *`PRF`* curves; they were assumed to be population-specific for *`PRF`*_*pop*_ and subject-specific for models *`PRF`*_*sbj,d*_ to *`PRF`*_*sbj,sdd*_. In these models, ***B*** (*β*_*c*_, *β*_*r*_) reflects the amount of explained variance of each physiological variable on the GS. The last 7 models (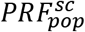 to *`PRF`*_*sc*_) are based on Eq. 6.

The 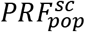 model assumes that ***G*** is population-specific, the models 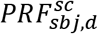 to 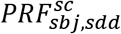 assume that ***G*** is subject-specific, while the *`PRF`*_*sc*_ model assumes that it is scan-specific. In the last seven models ***B*** (*β*_1,*c*_, *β*_2,*c*_, *β*_1,*r*_, *β*_2,*r*_) defines the amount of explained variance of the physiological variables on the GS as well as the shape of the *`PRF`* curves. As described in Section 2.4.1, in all 13 models, the interior-point gradient-based algorithm was applied after being initialized with the parameter values obtained from the population-specific model *`PRF`*_*pop*_ (for a pseudocode of the algorithm used to estimate the *`PRF`* parameters in all 13 models, please see Supplementary Table 2).

The following notation was adopted: the subscript in the six models *`PRF`*_*pop*_ to *`PRF`*_*sbj,sdd*_ (group of models II in Table 1) indicates whether ***G*** and ***R*** were estimated from the entire *population* (‘pop’) or a different scan of the same *subject* (‘sbj’). The subscript ‘sbj’ is always followed by a comma and a string consisting of the letters ‘d’ and ‘s’. The letters ‘d’ and ‘s’ in this string indicate whether parameter estimation and model performance assessment were implemented using data from one or more scans of a particular subject collected during a *different* or the *same* scanning session respectively. For example, *`PRF`*_*sbj,dd*_ denotes that ***G*** and ***R*** were estimated using data from two scans collected during one session (e.g. R1a/R1b) and model performance was assessed using data from a scan collected during a different session (e.g. R2a) from the same subject.

Similarly, the subscript in the six models 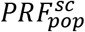 to 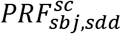 (group of models III in Table 1) indicates whether ***G*** was estimated from the entire *population* (‘pop’) or a different scan of the same *subject* (‘sbj’) with respect to the scan that the model performance was assessed on. In addition, the superscript ‘sc’ in the six models 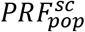 to 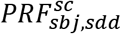 (Table 1) indicates that, even though ***G*** may be population- or subject-specific, the ultimate *`PRF`* shape is different for each *scan*. This is due to that ***B***, which was estimated for each scan separately, in the six models of group III consists of four parameters (i.e. *β*_1,*c*_, *β*_2,*c*_, *β*_1,*r*_, *β*_2,*r*_), in contrast to the 6 models of group II for which ***B*** consists of two parameters (i.e. *β*_*c*_, *β*_*r*_), allowing some flexibility in the shape of the *`PRF`*. The notation 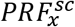 used for the models in group III indicates that, first, the parameter vector ***G*** was estimated from the dataset *x* (i.e. data from the same population or subject) and, subsequently, ***B*** was estimated from the same scan in which model performance was assessed. Finally, the subscript ‘sc’ for the last model (*`PRF`*_*sc*_) indicates that both ***G*** and ***B*** were estimated and validated from data collected during the same scan; therefore, this was the most flexible model.

Fig. 1 shows schematic examples of training and validation sets for each model. To assess the performance of the population-specific models *`PRF`*_*pop*_ and 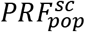, a leave-one-out cross-validation (LOOCV) approach was implemented at the first-level, whereby the *`PRF`* curves were obtained using training data from 40 subjects and validated with data from the remaining subject. In the case of subject-specific models of the form 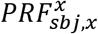, we considered all possible scan combinations (instead of using one of the scans as the validation data set and all remaining scans as the training set) to examine the effect of session on the variability of *`PRF`* curves (e.g. *`PRF`*_*sbj,d*_ vs *`PRF`*_*sbj,d*_) as well as the dependence of the model performance on the amount of training data (e.g. *`PRF`*_*sbj,d*_ vs *`PRF`*_*sbj,dd*_) in more detail.

**Fig. 1.**
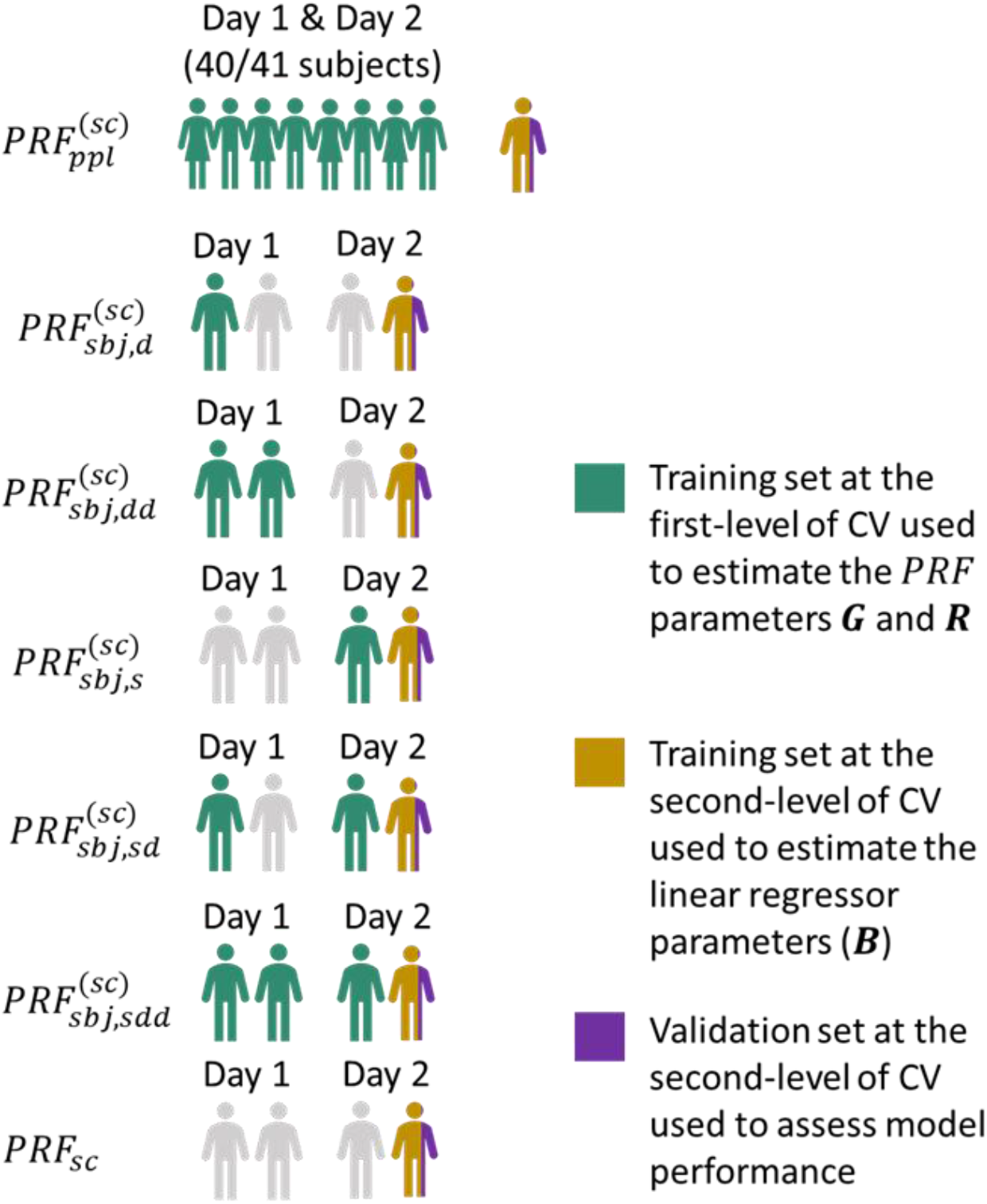
Schematic examples of training and validation sets for the two-level cross-validation (CV) framework used to compare the performance between different *`PRF`* models.

##### 2.4.2.2 Assessment of model performance

For a given scan and model, the *`PRF`* parameters (***G*** and ***R***) were extracted at the first-level as described earlier (Section 2.4.2.1). Subsequently, at the second-level, a 3-fold cross validation approach was implemented using the validation set of the first-level to prevent overfitting (Fig. 1). Each scan in the validation set of the first-level CV was partitioned into three segments of about 5 minutes each. One segment was used as the validation set at the second-level for assessing the performance of the model and the remaining two segments were used as the training dataset (at the second-level). This step was repeated three times with each of the three segments used exactly once as the validation data. In each fold, linear regression analysis was performed on the training set to estimate ***B*** (5^th^ column of Table 1). Subsequently, the estimated ***B*** was used on the validation set (second-level), and the correlation of the model prediction with the GS was calculated. Finally, the mean correlation across the three folds was calculated. For the *`PRF`*_*sc*_ model, one-level cross-validation was used. Specifically, the estimation of both ***G*** and ***B*** was performed on the training set of each fold (i.e. two segments of 5 minutes) and subsequently used on the validation dataset (i.e. the remaining segment of that scan).

For each model, the *`PRF`* parameters and performance assessment can be obtained from different scan combinations. For instance, in the case of *`PRF`*_*sbj,d*_ for a given subject, the *`PRF`* parameters can be estimated from scans R1a (or R1b) and the performance can be assessed on scans R2a (or R2b), yielding 8 total combinations. Therefore, for a given subject, the final performance of a model was assessed for all possible combinations and the average of the obtained values was used at the group level to compare the performance between all examined models (Table 1).

##### 2.4.2.3 Effect of sample size and duration of scan on PRF parameter estimation

To investigate the effect of the number of subjects on parameter estimation in population-specific models (*`PRF`*_*pop*_ and 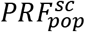), we repeated the assessment of model performance for the models *`PRF`*_*pop*_ and 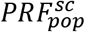 for the following subject numbers: 1-10, 15, 20, 25, 35 and 40. The subjects were randomly chosen and this part of the analysis was repeated ten times to eliminate biases from “representative” or “non-representative” subjects chosen in a particular iteration. In each iteration, subjects that were not chosen for parameter estimation were included in the validation set. Similarly, to examine the effect of scan duration on the estimation of *`PRF`* parameters and investigate the minimum required duration in scan-specific models, we evaluated the performance of the models *`PRF`*_*pop*_ and *`PRF`*_*sc*_ for scan durations between 1 to 15 minutes in steps of one minute. The assessment of the model performance was done using a 3-fold cross validation approach as described in Session 2.4.2.2.

##### 2.4.2.4 Performance of *PRF* estimation on long TR data

This study employs a novel methodological framework for extracting *`PRF`* curves that allows the convolution of physiological variables (HR and RF) with the *`PRF`* curves to be done in a pseudo-continuous time-domain at a 10 Hz sampling frequency regardless of the TR in the fMRI data. Earlier studies examining *`PRF`* curves downsampled the physiological variables to the fMRI sampling rate before convolution (or deconvolution), which may smooth out the effect of high-frequency physiological fluctuations of HR and RF (or RVT). On the other hand, our approach avoids this possible loss of high-frequency information in HR and RF, and therefore can allow the estimation of curves with faster dynamics than what has been previously reported. However, earlier studies that examined *`PRF`* curves have used datasets with longer TR in fMRI pulse sequence compared to the one examined in this study (TR=0.72 s). To examine whether the method for `PRF` estimation proposed here yields similar results for longer TRs in terms of the shape of the *`PRF`* curves, we artificially reduced the sampling rate of the fMRI data by a factor of 5, obtaining a similar TR (3.6 s) with previous studies (Birn et al., 2008; Chang et al., 2009), and repeated the estimation of the *`PRF`*_*sc*_ curves in both datasets (short and long TR), omitting the cross-validation analysis.

##### 2.4.2.5 Comparison of population-, scan- and voxel-specific *PRF* curves in individual voxels

Here, we aimed to examine the variability of the *`PRF`* curves across voxels. To this end, we compared the performance of a subset of models considered in 2.4.2, particularly the standard models *`PRF`*__*stand*__, the *`PRF`*_*pop*_, 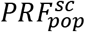 and *`PRF`*_*sc*_ models, in individual voxels. In addition, we examined the performance of a voxel-specific *`PRF`* model, termed 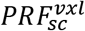. 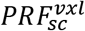 was simply an extension of *`PRF`*_*sc*_ that allowed variability in the shape of the *`PRF`* curve across voxels. The *`PRF`* curves for the first four models (standard models *`PRF`*__*stand*__, *`PRF`*_*pop*_, 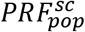 and *`PRF`*_*sc*_) were estimated based on the GS of one or more scans as described earlier in Section 2.4.2. These curves were subsequently used to extract two physiological regressors that were included in the GLM for a voxel-wise analysis. On the other hand, for the 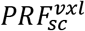 model, the four gamma functions corresponding to parameter vectors ***G*** and ***B*** of *`PRF`*_*sc*_ were used to extract four physiological regressors for the GLM. Due to that 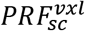 had more physiological regressors in the GLM than the other models examined in this section (4 vs 2), a 3-fold cross-validation approach was implemented to assess model performance as described in Section 2.4.2.1.

As the distinction of the five different models is in some cases subtle, we refer the readers to Supplementary Fig. 3 for a diagram that illustrates the steps involved in estimating each model. Specifically, the diagram shows how the physiological regressors needed for the voxel-wise GLM analysis are extracted in practice. Regarding the notation adopted in this work, the subscript always indicates the level at which the parameter vector ***G*** was estimated from (scan-, subject- or population-level). In the cases that the final shape of the *`PRF`* curves was defined at a lower level (voxel-, scan- or subject-level) compared to ***G***, the lower level is indicated by the superscript. For example, 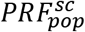 indicates that ***G*** was estimated from the population, while the final shape of the *`PRF`* curve was defined at the scan level using the GS as the fitting target. Similarly, 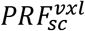 indicates that ***G*** was estimated at the scan level using the GS and the final *`PRF`* curve was defined at the voxel level. Note that the 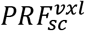 curves were implicitly defined for each voxel when the parameter vector ***B*** of each voxel was estimated using the voxel-wise GLM.

The comparison of the models was restricted to regions of interest (ROIs) where the models explained significant variance. Specifically, these ROIs were defined for each model separately and included the 5% of voxels in the brain with the highest correlation between the voxel time-series and the prediction of the corresponding model. The aforementioned five *`PRF`* models were examined for *`CRF`* and *RRF* separately as well as both *`CRF`* and *RRF*, yielding 15 models in total. The comparison between the 15 models was repeated on FIX-denoised data using ROIs for each model the ones derived from the original data. The ROIs were defined for each model and scan separately, possibly resulting in different voxel sets examined each time. To examine whether this approach may somehow bias the results, we repeated the analysis using a fixed ROI. This ROI was defined as the voxels in the brain with the 5% highest correlation values in the group-level map obtained using the standard models *`PRF`*__*stand*__. Finally, to examine the effect of spatial smoothing on the performance of *`PRF`* models, the comparison between the 15 models on the raw data was performed with a FWHM value of 0 mm and 6 mm in addition to the value of 3 mm used for the main analysis.

For visualization purposes, the statistical maps shown here were overlaid on structural images after being transformed to structural space with FSL’s FLIRT registration tool (Jenkinson and Smith, 2001) as incorporated in the MANGO software (Lancaster, Martinez; www.ric.uthscsa.edu/mango).

#### 2.4.4 Comparison of PRF curves for different variants of GS

While in the previous sections we considered the mean time-series from voxels in the whole brain (WB) to obtain the GS, an important question is whether other variants of the GS are more sensitive to fluctuations in HR and RF. To this end, we repeated the *`PRF`* estimation for the model that was found to yield the best performance (i.e. *`PRF`*_*sc*_), considering as GS the mean time-series derived from the following ROIs: 1. WB, 2. gray matter (GM), 3. white matter (WM) and 4. the cerebrospinal fluid (CSF) compartment. In addition, we examined whether preprocessing of the fMRI data before *`PRF`* parameter estimation significantly affects model performance by using different GS variants from each ROI before and after fMRI data preprocessing (i.e. regressing out the motion realignment parameters and high-frequency cardiac and respiratory regressors estimated with RETROICOR). The variants of the GS derived from the preprocessed fMRI data are denoted later in the text with the subscript clean (e.g. WB_clean_) to distinguish them from those derived from the raw fMRI data.

## 3. Results

### 3.1 Variability in physiological measurements

The mean HR and BR values during resting conditions exhibited considerable variability across the 164 scans, with the mean HR and BR ranging between 46-106 bpm and 13-23 rpm, respectively. To examine the variability across subjects and sessions, all scans were grouped in pairs a) from different subjects (*N* = 13120; for a pair of scans the first scan was picked from all 164 scans and the second scan from the 160 scans of the remaining 40 subjects. Thus, the number of unique pairs was 164 x 160 divided by two as the order does not matter), b) from different sessions of the same subject (*N* = 41 ∙ 2 ∙ 2 = 164), and c) from scans of the same session (*N* = 41 ∙ 2 = 82). The mean HR differences obtained from the scan pairs in groups a, b and c yielded standard deviation values of 16, 8 and 2 bpm respectively, and the corresponding variance values were found to be significantly different (*F*-test; *p*-value: <10^−49^). Similarly, the differences in BR obtained from the scan pairs in groups a, b and c yielded standard deviation values of 3.1, 1.7 and 1.0 rpm, and the corresponding variance values were found again to be significantly different (*F*-test; *p*-value: <10^−15^). In other words, the between subject variability in mean HR and BR was significantly larger compared to the within-subject variability and, in turn, the between session within-subject variability was larger than the within session variability. In Fig. 2, we show the variability of mean HR and BR across subjects, sessions and scans for 10 representative subjects, which illustrates the aforementioned statistical differences. For instance, significant differences were found between scans of different subjects such as in the case of subjects S406432 (light blue color) and S555348 (brown color), whereby all four scans of the former subject are characterized by lower mean HR and BR compared to the four scans of the latter one. Also, significant variability is found between sessions within-subject such as in the case of subject S203923 (cyan color) whereby both mean HR and BR increase from the first to the second session.

**Fig. 2.**
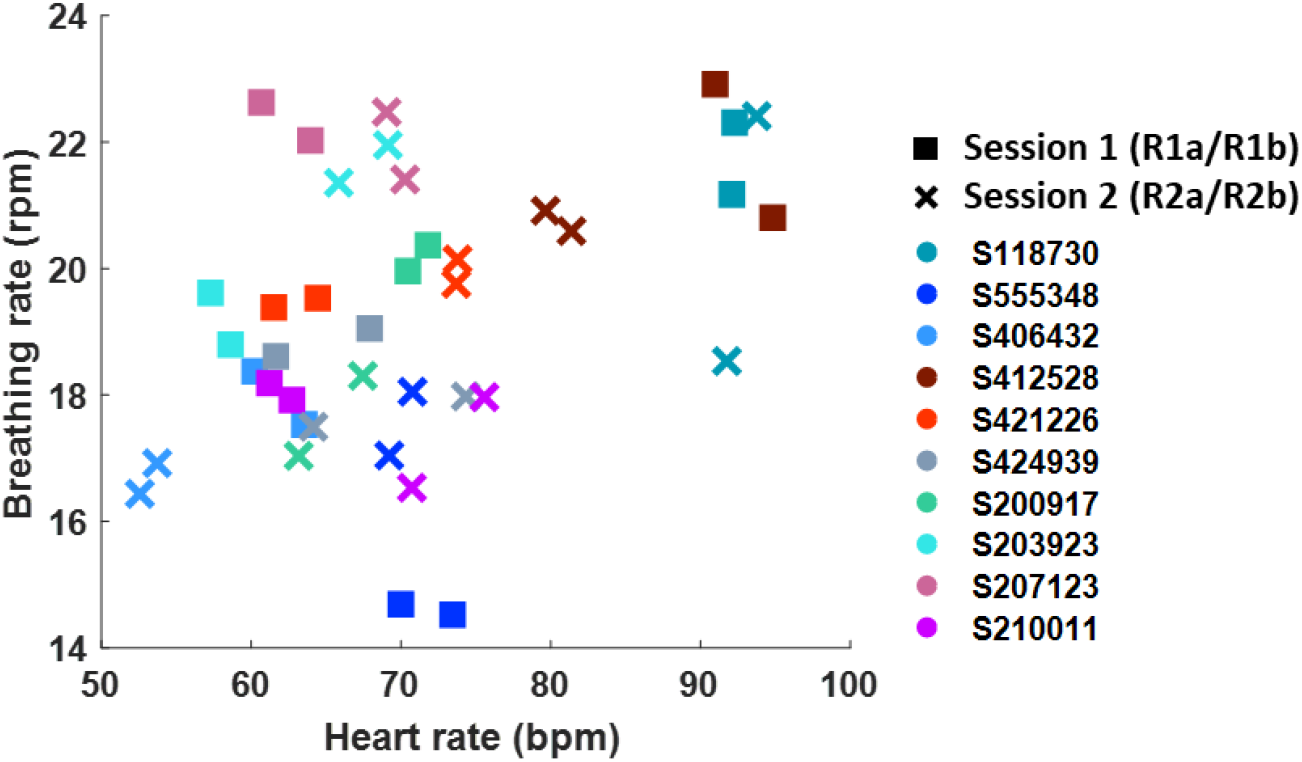
Scatterplot of mean heart rate (HR) and breathing rate (BR) from scans of 10 representative subjects. Squares and crosses correspond to scans from sessions (days) 1 and 2, respectively, whereas each color indicates a different subject. Both mean HR and BR vary less across scans of the same session than scans across different sessions within-subject. In turn, mean HR and BR across the four scans of each subject exhibited lower variability compared to scans from different subjects.

### 3.2 Variability in the shape of *PRF* curves across scans

This section examines the variability in the shape of the *`PRF`*_*pop*_, 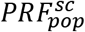 and *`PRF`*_*sc*_ curves. For all 164 scans examined in this study, it was found that the GS was strongly correlated to cardiac and breathing activity. The first column of Fig. 3 shows the optimal *`CRF`*_*pop*_ and *RRF*_*pop*_ curves estimated for the 41 subjects using the *`PRF`*_*pop*_ model that assumes the same *`CRF`* and *RRF* for the entire population. Both the *`PRF`*__*stand*__ and *`PRF`*_*pop*_ curves have a bimodal shape with a positive peak followed by a negative one. However, the amplitude and time of the peaks differ between the curves. The peaks in the *`CRF`*_*pop*_ appear at much earlier time lags compared to *`CRF`*__*stand*__ (Chang et al., 2009; 1.2 and 7.0 s for *`CRF`*_*pop*_, vs. 4.1 and 12.4 s for *`CRF`*__*stand*__). Faster dynamics were also observed in the estimated *RRF*_*pop*_ compared to *RRF*__*stand*__ (Birn et al., 2008). Specifically, the positive and negative peaks of the *RRF*_*pop*_ were found to be located at 2.0 and 12.8 s, respectively, whereas the corresponding peaks in the *RRF*__*stand*__ are at 3.1 and 15.5 s. Moreover, the estimated *`PRF`*_*pop*_ curves return to baseline faster than the corresponding *`PRF`*__*stand*__ curves. Finally, *`CRF`*_*pop*_ and *RRF*_*pop*_ exhibited a stronger positive and negative peak respectively compared to the standard curves (factor of ~2).

**Fig. 3.**
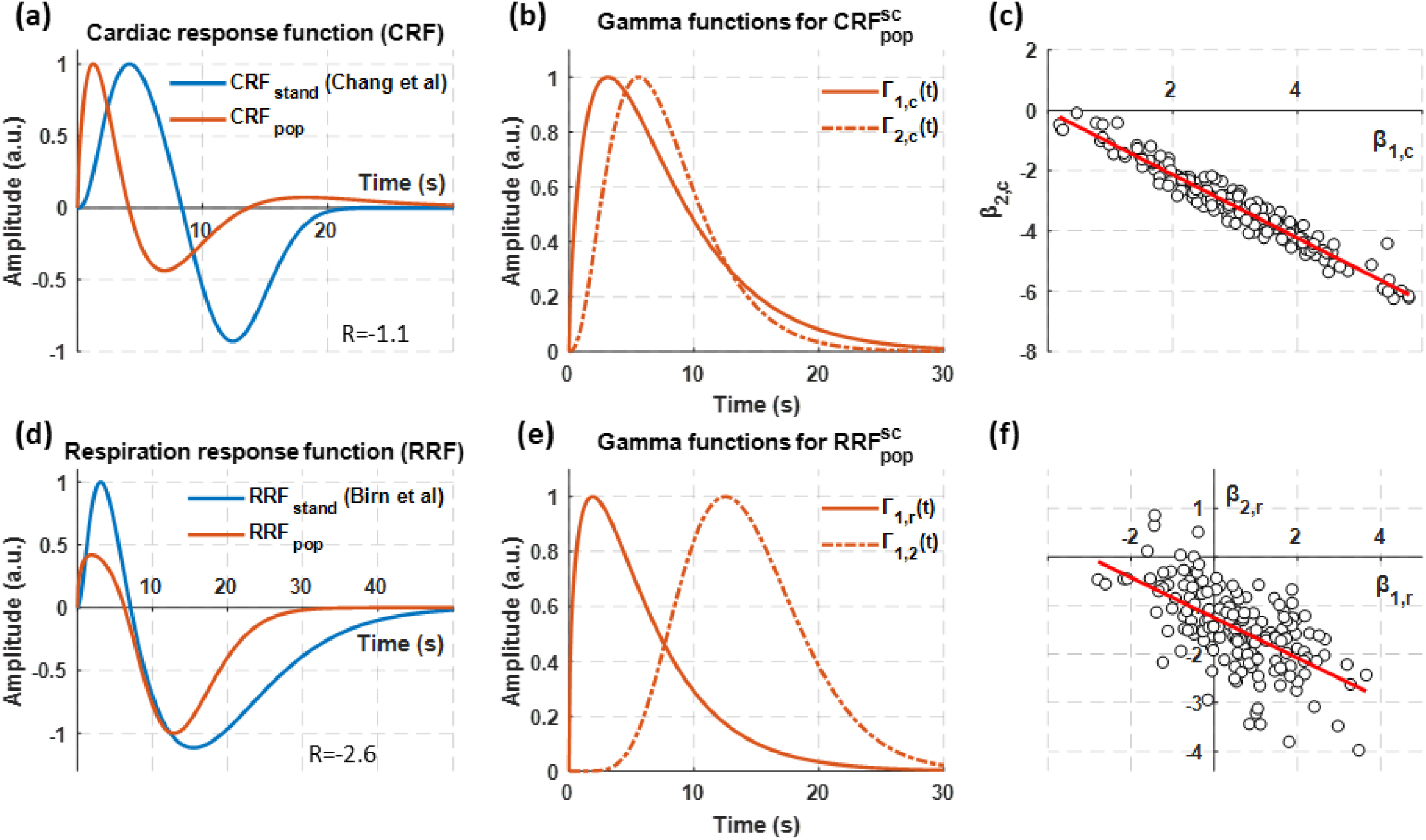
Estimated gamma functions for the 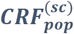 and 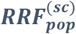 curves. (a) Standard *`CRF`*__*stand*__ and estimated *`CRF`*_*pop*_ curves. (b) The two gamma functions used to construct the population-specific *`CRF`*_*pop*_ and 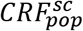 curves. (c) Scatterplot of parameters in ***B*** for the 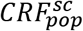 model. The circles correspond to the 164 scans and the values of *β*_1,*c*_ and *β*_2,*c*_ reflect the amplitude of the gamma functions *Γ*_1,*c*_ and *Γ*_2,*c*_ shown in (b). In a similar manner, (d), (e) and (f) illustrate the results for the models related to *RRF*.

The second column of Fig. 3 shows the gamma functions that were used to construct the *`PRF`*_*pop*_ and 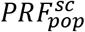 curves while the third column presents scatterplots of ***B*** (*β*_*c*_1, *β*_*c*_2, *β*_*r*_1, *β*_*r*_2), which defined the shape of the 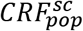 and 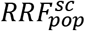 curves, for all scans. Specifically, *β*_*c*_1 (*β*_*r*_1) and *β*_*c*_2 (*β*_*r*_2) correspond to the sign and magnitude of the first and second peak of the 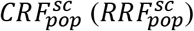 curve, respectively. The parameters of the four gamma functions are listed in Table 2. All parameters in ***B*** for 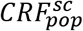 are located on the 4^th^ quadrant (Fig. 3c) and can be approximated well with a straight line that crosses the origin of the plane with a slope R of −1.1. However, the small deviation of each circle from the straight line suggests that the 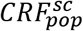 curves had small differences compared to the *`CRF`*_*pop*_ curve shown in Fig. 3b. Furthermore, the parameters in ***B*** for the 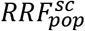 curves (Fig. 3f) indicate an even larger variability of the curves across scans compared to the *RRF*_*pop*_ shown in Fig. 3e.

**Table 2.** Parameters for the 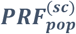 model. The parameter vectors ***τ*** and ***δ*** used in Eq. 5 are an approximate measure for the time of peak and dispersion of the gamma function. The full width at half maximum (FWHM) was calculated numerically.

| | $CRF_{pop}^{(sc)}$ | | $RRF_{pop}^{(sc)}$ | |
| --- | --- | --- | --- | --- |
| | $\Gamma_{c_1}(t)$ | $\Gamma_{c_2}(t)$ | $\Gamma_{r_1}(t)$ | $\Gamma_{r_2}(t)$ |
| Time to peak $\tau$ (s) | 3.1 | 5.6 | 1.9 | 12.5 |
| Dispersion $\delta$ (s) | 2.5 | 0.9 | 2.9 | 0.5 |
| FWHM (s) | 9.2 | 8.3 | 7.0 | 11.1 |
| Ratio $R$ ($\beta_2/\beta_1$) (only for $PRF_{pop}$) | N/A | -1.1 | N/A | -2.6 |

Fig. 4 illustrates the estimated *`PRF`*_*sc*_ curves for two subjects that demonstrated strong association between the GS with both HR and breathing pattern. It also shows the physiological variables HR and RF, as well as the estimated regressors that maximized the correlation with the GS. For the majority of the examined subjects, including the two subjects whose results are shown in Fig. 4, it was observed that HR explains faster fluctuations of the GS compared to RF. Additionally, we observe that even though the two subjects had almost the same mean HR, the corresponding dynamic patterns were very different. Specifically, the HR of subject 210415 was relatively stable with sporadic abrupt increases whereas the HR of subject 30717 exhibited faster fluctuations. With respect to the estimated *`CRF`*_*sc*_ curves, subject 210415 was characterized by a more abrupt increase and faster return to baseline. The RF time-series of these two subjects also exhibited different profiles, while their *RRF*_*sc*_ curves differed significantly from the canonical *RRF*__*stand*__ and the population-specific *RRF*_*pop*_ curve. The rest of the three scans of subject 210415 exhibited *RRF*_*sc*_ curves similar to the *RRF*_*pop*_ (Fig. 3d) while the curves found for the rest of the three scans of subject 307127 were similar to the curve derived from the R2a scan shown in Fig. 4 (Supplementary Figs. 4-7). For visual inspection of the model fit for all scans please see our fishare repository (https://doi.org/10.6084/m9.figshare.c.4585223.v4; Kassinopoulos and Mitsis, 2019).

**Fig. 4.**
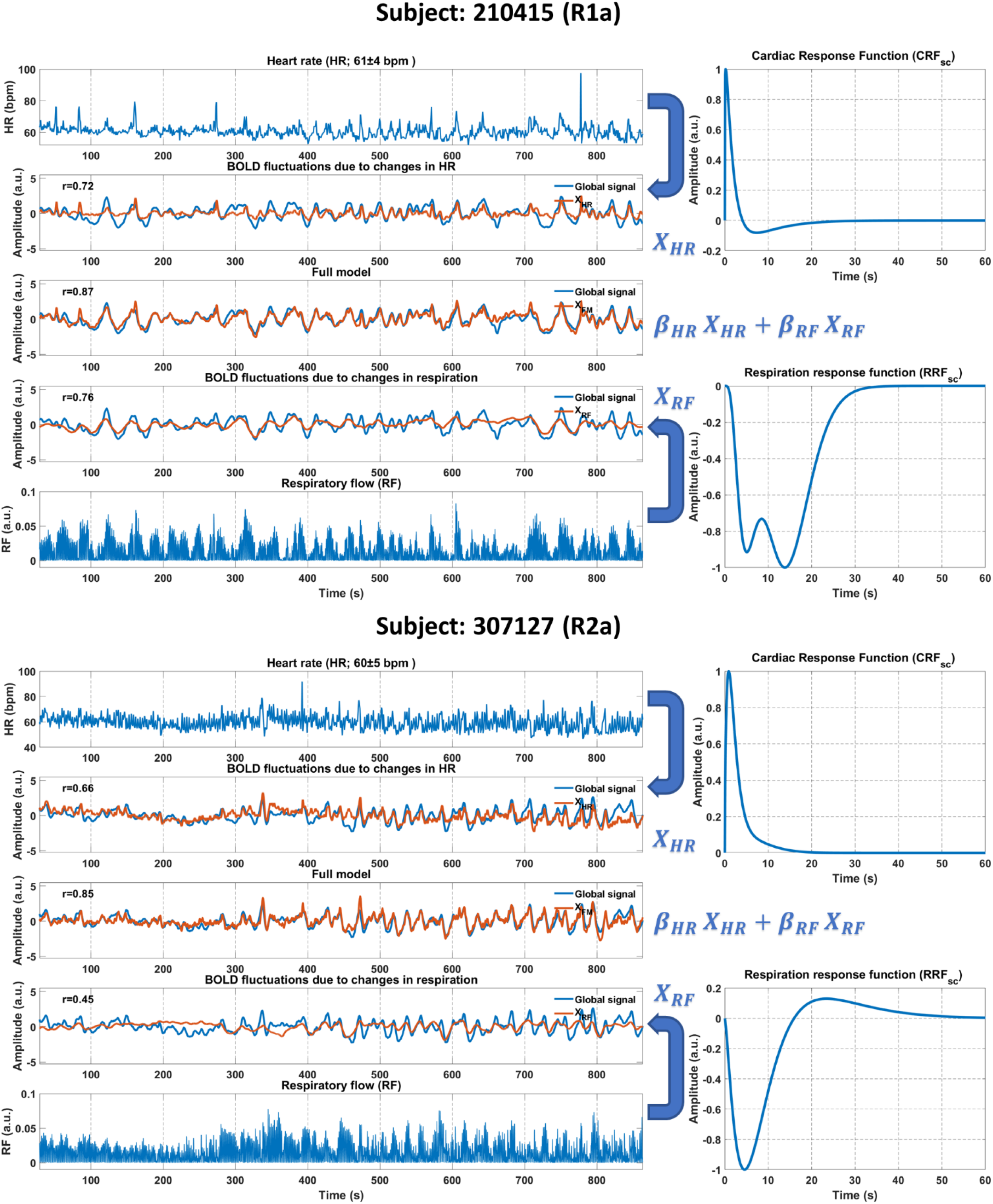
Demonstration of the *PRF*_*sc*_ model performance for two subjects. The HR (1^st^ row of each subject’s panel) and RF (5^th^ row) time-series were derived from the recorded physiological signals (photoplethysmograph and respiratory belt, respectively). Subsequently, the scan-specific curves *`CRF`*_*sc*_ and *RRF*_*sc*_ were obtained after estimating the corresponding *`PRF`* parameters (right column). The physiological regressors *X*_*HR*_ and *X*_*RF*_ shown in the 2^nd^ and 4^th^ row respectively were obtained as the convolution between HR/RF with *`CRF`*_*sc*_/*RRF*_*sc*_. The parameters *β*_*HR*_ and *β*_*RF*_ were estimated by maximizing the variance of the GS explained by the model (3^rd^ row).

To better understand the properties of the scan-specific *`PRF`*_*sc*_ curves, we investigated whether the times of positive and negative peak depend on physiological variables (e.g. mean and variance of HR). Among the different combinations that we tested, significant correlations were found only between the shape of the *`CRF`*_*sc*_ curves and the subjects’ HR. Specifically, shorter times for the negative *`CRF`*_*sc*_ peak correlated with higher mean HR values (Fig. 5a). Furthermore, we examined whether HR and RF fluctuations had a stronger effect on the GS under specific physiological states. Fig. 5b shows that HR values were strongly negatively correlated with the fraction of the GS explained by the cardiac regressor (as quantified by the correlation coefficient between model prediction and GS), whereas RF variance values were found to be significantly correlated to the fraction of the GS explained by the respiratory regressor (Fig. 5c).

**Fig. 5.**
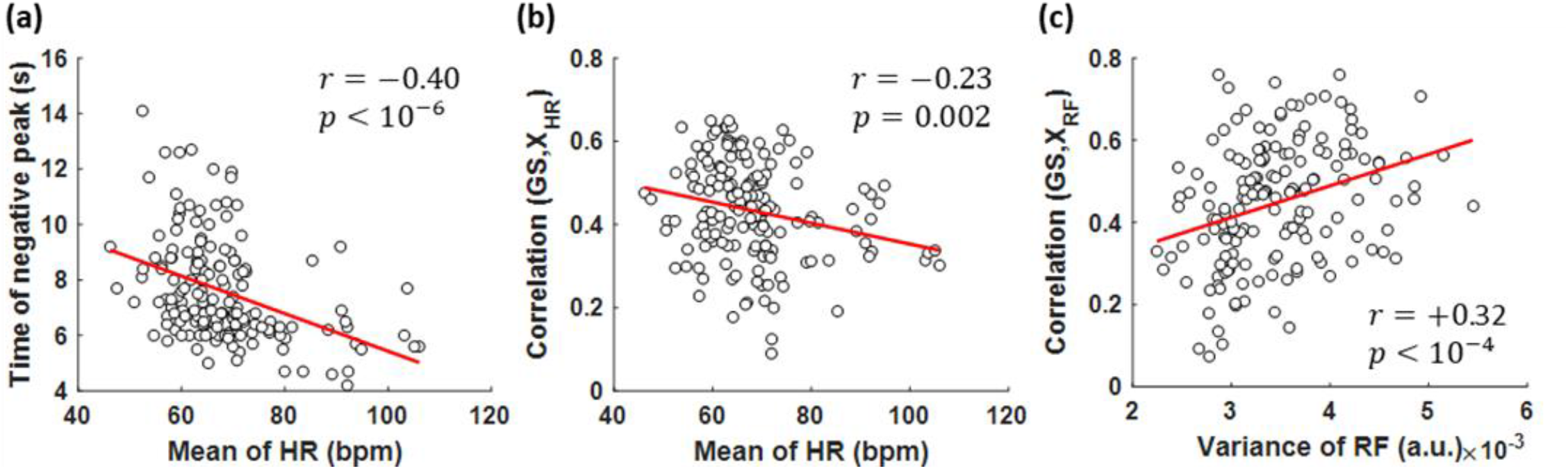
Scatterplots of physiological variables and features from the *PRF*_*sc*_ models. (a) The time-to-peak values for the negative *`CRF`*_*sc*_ peak were negatively correlated with mean HR. (b) The mean HR was negatively correlated with the correlation coefficient between the cardiac regressor *X*_*HR*_ and GS. (c) The variance of RF was strongly correlated with the correlation coefficient between the respiratory regressor *X*_*RF*_ and GS.

#### 3.3 Comparison of population-, subject-, session- and scan-specific *PRF* curves

The population-specific curves *`PRF`*_*pop*_ yielded a significant increase in the mean correlation between the corresponding physiological regressors and the GS compared to the standard curves *`PRF`*__*stand*__ (from 0.30 to 0.51; Fig. 6, *p*<0.0001 uncorrected). Gradually making the `PRF` curves more scan-specific yielded additional improvements in the model performance. Specifically, 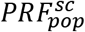 (whereby ***B*** (i.e. *β*_1,*c*_, *β*_2,*c*_, *β*_1,*r*_, *β*_2,*r*_) that determines the shape of the *`PRF`* curves was allowed to vary across scans) yielded a significant increase in the mean correlation compared to *`PRF`*_*pop*_ (0.53; *p*<0.01 – Fig. 6), while optimizing all the parameters for each scan separately (*`PRF`*_*sc*_) further increased the mean correlation to 0.56 (p<0.0001 compared to 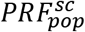 – Fig. 6). These increases were not due to overfitting the data, as we assessed model performance using different training and testing datasets.

**Fig. 6.**
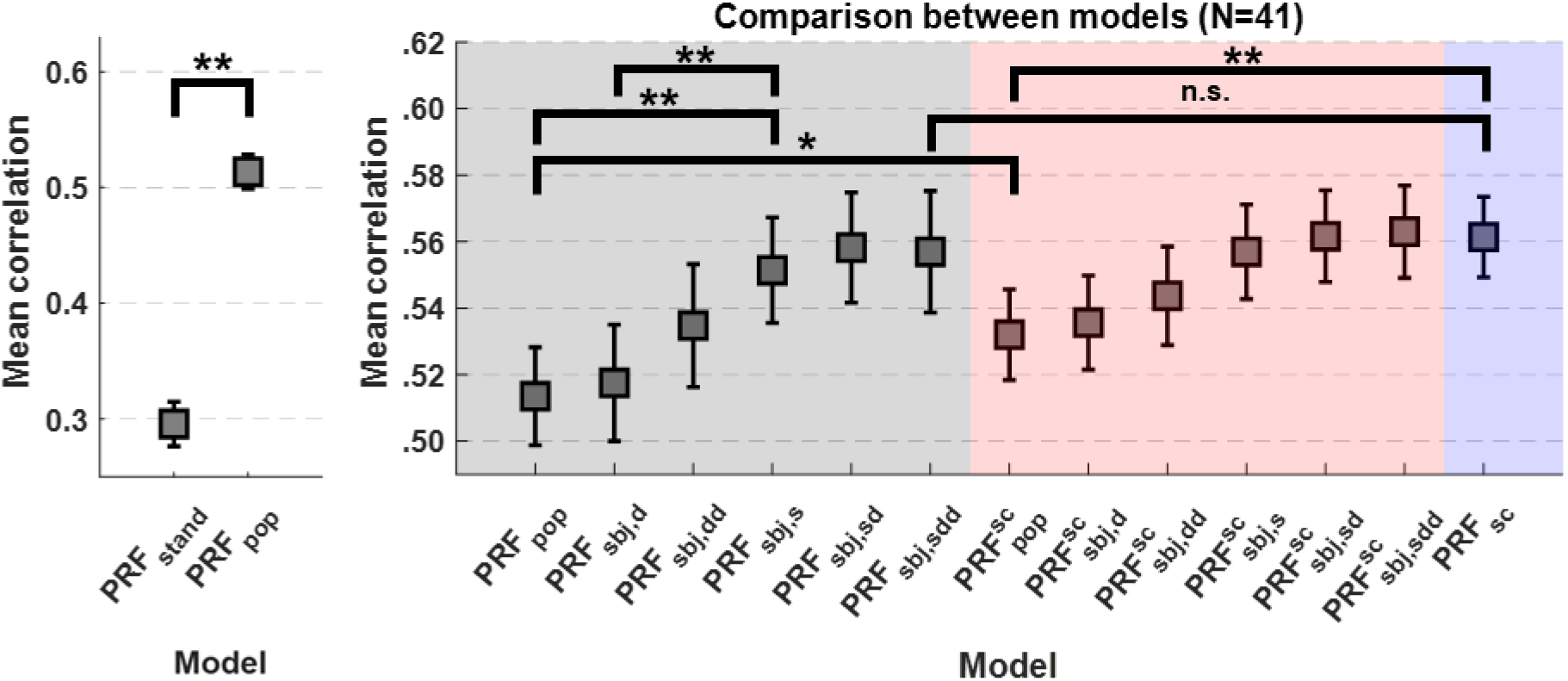
Mean correlation between the prediction of each model with the GS. The squares and error bars indicate the mean and standard error of the means across all subjects. The different colors in the shaded areas correspond to the groups of models II, III and IV from Table 1. The population-specific model *`PRF`*_*pop*_ yielded a significant mean correlation increase compared to the standard curves (*`PRF`*__*stand*__). However, models with more flexibility yielded further improvements in performance, suggesting that *`PRF`* curves vary both from subject to subject as well as between scans within a subject. \**p* < 0.01; \*\**p* < 0.0001.

As illustrated in Fig. 6, subject-specific models trained and assessed using different sessions (*`PRF`*_*sbj,d*_) yielded slightly higher mean correlation values compared to the population-specific model *`PRF`*_*pop*_, which were however not statistically significant. Increasing the amount of data used for training (two scans of 15-minute duration from one session; *`PRF`*_*sbj,dd*_) further improved performance, but this improvement was again non-significant compared to *`PRF`*_*pop*_. On the other hand, a significantly improved fit was achieved when the models were estimated and validated from different scans collected during the same session (*`PRF`*_*sbj,d*_). Importantly, the *`PRF`*_*sc*_ model, whereby parameter estimation and assessment of performance was done by splitting the same scan performed equally well with *`PRF`*_*sbj,d*_ but also with *`PRF`*_*sbj,sdd*_ and 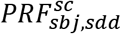, which used the maximum number (3) of within-subject scans for training. This finding suggests that estimating subject-specific *`PRF`* curves from data collected on different sessions or scans, regardless of the amount of data, does not offer any benefit compared to scan-specific *`PRF`*_*sc*_ curves derived from the same data.

#### 3.3.1 Effect of sample size and duration of scan on PRF parameter estimation

To assess the effect of the number of subjects on the reliability of the obtained population-specific models, we repeated estimation of the model parameters using 10 different randomized subject cohorts of different sizes (see Methods). In Fig. 7a, it can be seen that the performance of the population-specific models increased monotonically between 1 and 10 subjects and reached a plateau at a mean correlation of around 0.51 and 0.53 for the *`PRF`*_*pop*_ and 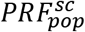 models, respectively. We also examined the effect of the scan duration, which may considerably affect model reliability, particularly in the case of *`PRF`*_*sc*_. Fig. 7b shows a comparison of the performance achieved for the least (*`PRF`*_*pop*_) and most (*`PRF`*_*sc*_) flexible models as a function of scan duration. The *`PRF`*_*pop*_ model yielded higher correlation values for short scan durations (0.44 for a one-minute duration), while these values stabilized around 0.51 for scan durations above 5 minutes. The *`PRF`*_*sc*_ model yielded poorer performance for scan durations shorter than 5 minutes, and its performance improved for longer scan durations, eventually reaching a mean correlation of 0.56 (15-minute scan duration). Interestingly, these two models yielded very similar performance for a 5-minute scan duration, which is the minimum scan duration typically used in rs-fMRI studies.

**Fig. 7.**
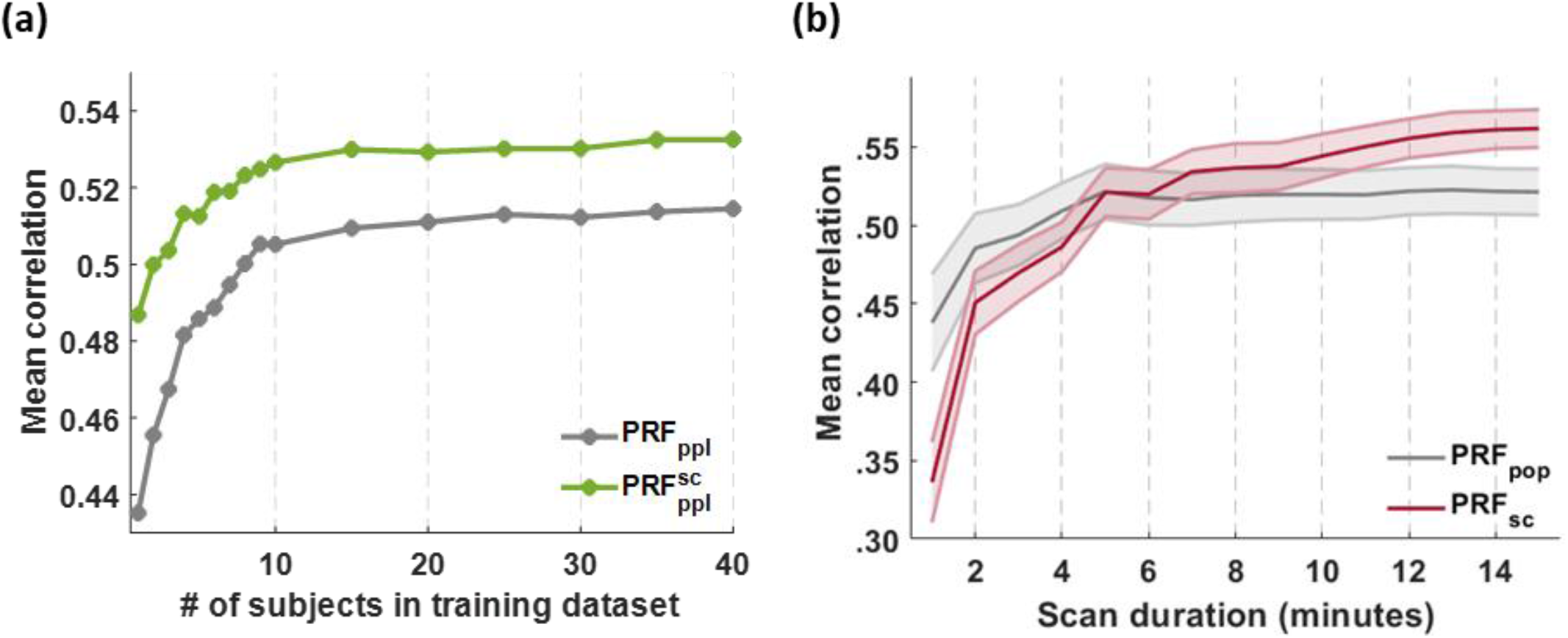
Effect of sample size and scan duration on the performance of *PRF* models. (a) The *`PRF`* parameters in the population-specific models *`PRF`*_*pop*_ and 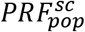 were estimated using subsets of subjects and the performance of the models was assessed on the remaining subjects. Both models reached a plateau at a sample size of around 10 subjects (b) The performance of the least flexible model (i.e. *`PRF`*_*pop*_) and the most flexible model (i.e. *`PRF`*_*sc*_) was assessed on reduced scan durations. Our results suggest that for scan duration longer than 5 minutes the scan-specific *`PRF`*_*sc*_ model explained more variance in GS than the *`PRF`*_*pop*_ model and a more profound difference was observed as the scan duration increased.

#### 3.3.2 Performance of PRF estimation on long TR data

Earlier in the paper it was shown that the *`CRF`*_*pop*_ curve is characterized by significantly faster dynamics than the standard curve *`CRF`*__*stand*__ reported in Chang et al. (2009). To examine whether this is due to the faster TR (i.e. 0.72 s) of the dataset examined here, we repeated the estimation of the *`PRF`*_*sc*_ curves with the same dataset downsampled to a TR of 3.6 s. Fig. 8 shows that the GS variance explained by HR and RF across scans is similar for both TRs. In addition, regarding the *`CRF`*_*sc*_ curves, as can be seen in Fig. 8 by the time of positive and negative peaks of the curves across scans, the short and long TR data yielded similar shapes. With respect to the *RRF*_*sc*_ curves, while we observe in Fig. 8 that the time of positive peak varied between the two TRs, visual inspection showed that the curves were again very similar. The scans that had large differences in the positive peak between the short and long TR, demonstrated also low amplitude in the positive peak. Therefore, the overall shape of the curves was essentially defined by the negative peak which was consistent between the two TRs.

**Fig. 8.**
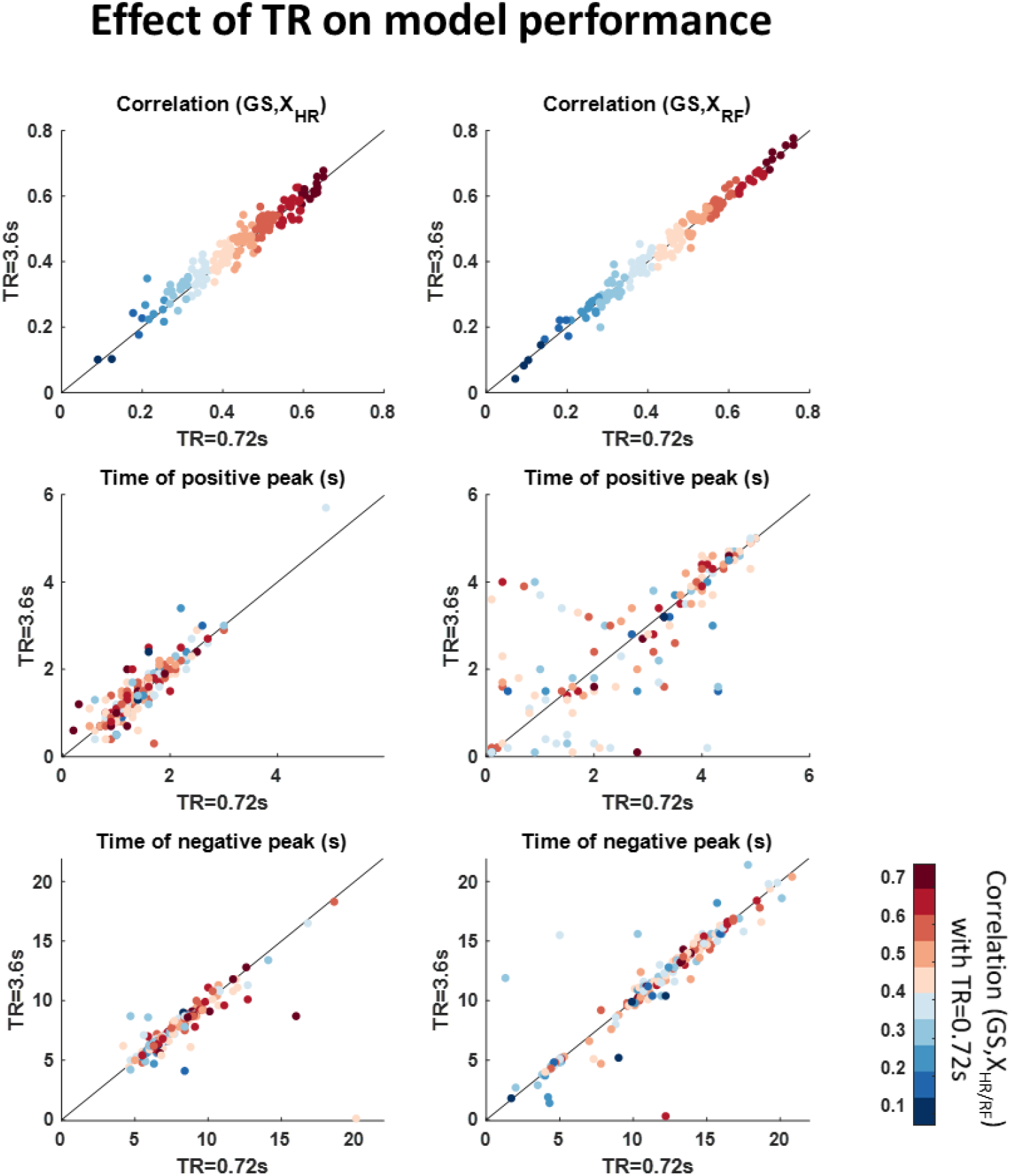
Effect of TR on model performance and characteristics of scan-specific curves *PRF*_*sc*_. The circles correspond to the 164 scans whereas each color in the left and right column indicates the fraction of GS variance explained with HR and RF, respectively, on the short TR data. Overall, the longer TR (3.6 s) yielded similar performance with the original TR (0.72 s) with respect to the variance explained and the shape of the *`PRF`*_*sc*_ curves. Even though the time of positive peak for *RRF*_*sc*_ exhibited differences between the two sampling rates, visual inspection showed that this is related to the small amplitude of the positive peak which causes the time of peak to vary without considerably affecting the overall shape.

### 3.4 Model performance in individual voxels

The comparison of the models was repeated on a voxel-wise basis in ROIs. The ROIs were defined for each model separately and included the 5% of voxels in the brain with the highest goodness of fit (see Methods). The analysis at the voxel-level yielded similar findings with the analysis based on the fit to the GS (Fig. 9). The full *`PRF`*_*sc*_ model (i.e. *`CRF`*_*sc*_ and *RRF*_*sc*_) yielded the best performance among all models with a mean correlation around 0.24 (note the lower values compared to GS). Importantly, the 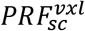 model demonstrated slightly lower mean correlation compared to the *`PRF`*_*sc*_ model, even though it allows variability in the estimated curves across voxels. For all the examined models, the breathing-related component exhibited higher mean correlation compared to the cardiac-related component, although the difference was not statistically significant (*p* > 0.05). The analysis for assessing model performance at the voxel level was also performed on resting-state fMRI data that were corrected for physiological noise with FIX. The results were similar to the results derived from the raw data, although the overall mean correlation was decreased in the latter case. The *`PRF`*_*sc*_ model again illustrated the best performance with a mean correlation of around 0.17. As the ROIs varied across models and scans, which implies that the models were compared on ROIs that were not entirely the same, we repeated the analysis using a fixed ROI based on the group-level map for the standard models *`PRF`*__*stand*__ (see Methods). Using a fixed ROI yielded similar results, even though there was an overall decrease in the obtained mean correlation for all models (Supplementary Fig. 8).

**Fig. 9.**
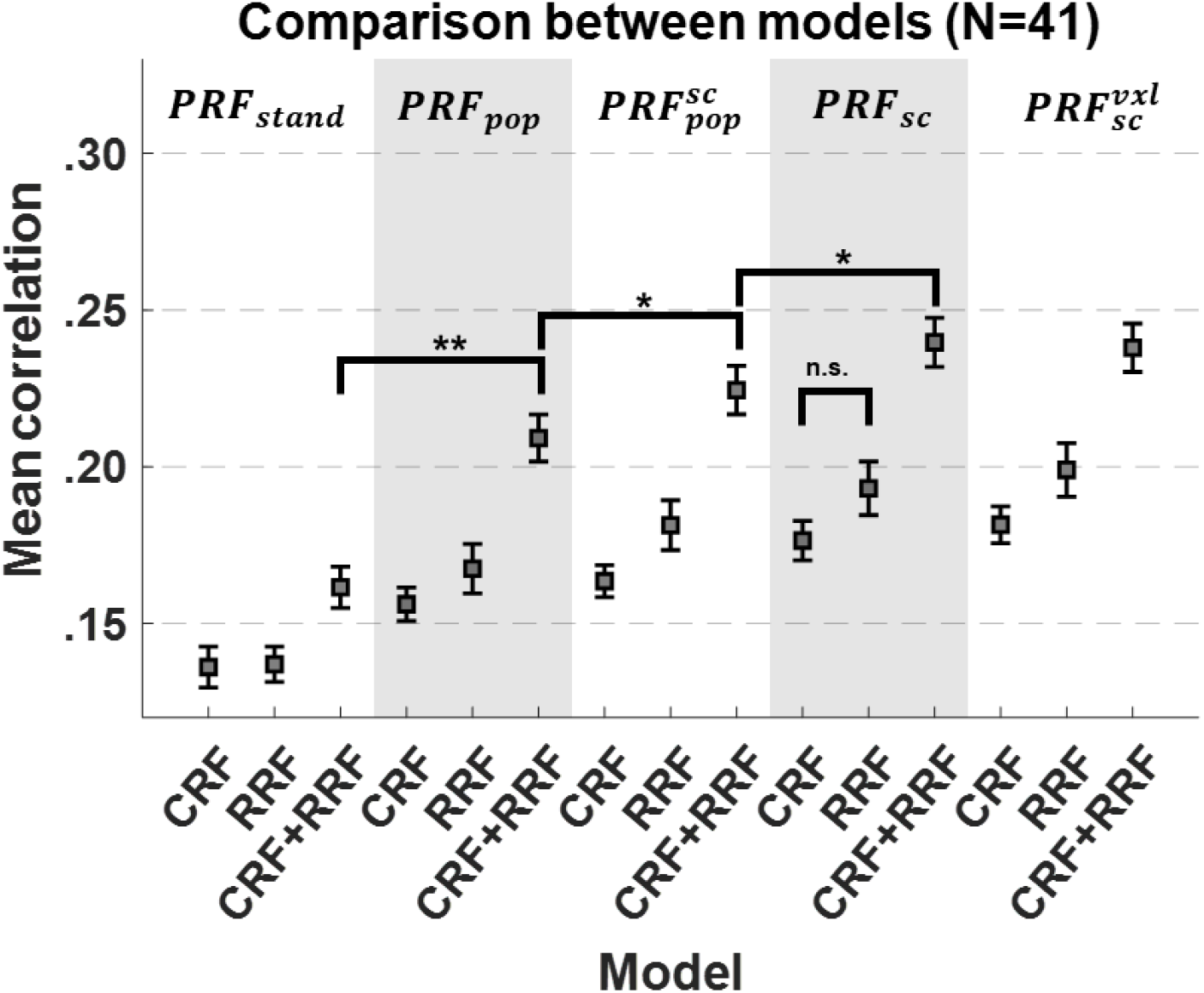
Correlation values between physiological model predictions and voxel-specific timeseries, averaged over all voxels within the ROI of each model. The squares and error bars indicate the mean and standard error of the means of all subjects. As in the case of GS-based analysis, the population-specific *`PRF`*_*pop*_ model yielded significantly increased performance compared to the standard methods (*`PRF`*__*stand*__). Overall, the best performance was achieved with the scan-specific model *`PRF`*_*sc*_. \**p* < 10^−8^; ** *p* < 10^−13^.

The brain areas affected by fluctuations in HR and breathing pattern were mainly areas in gray matter and close to blood vessels (Fig. 10). Unsurprisingly, the standard methods (*`PRF`*__*stand*__) and the proposed methods yielded similar maps regarding the areas more affected by physiological noise. However, the explained variance obtained using the proposed *`PRF`* curves was significantly higher compared to the standard curves.

**Fig. 10.**
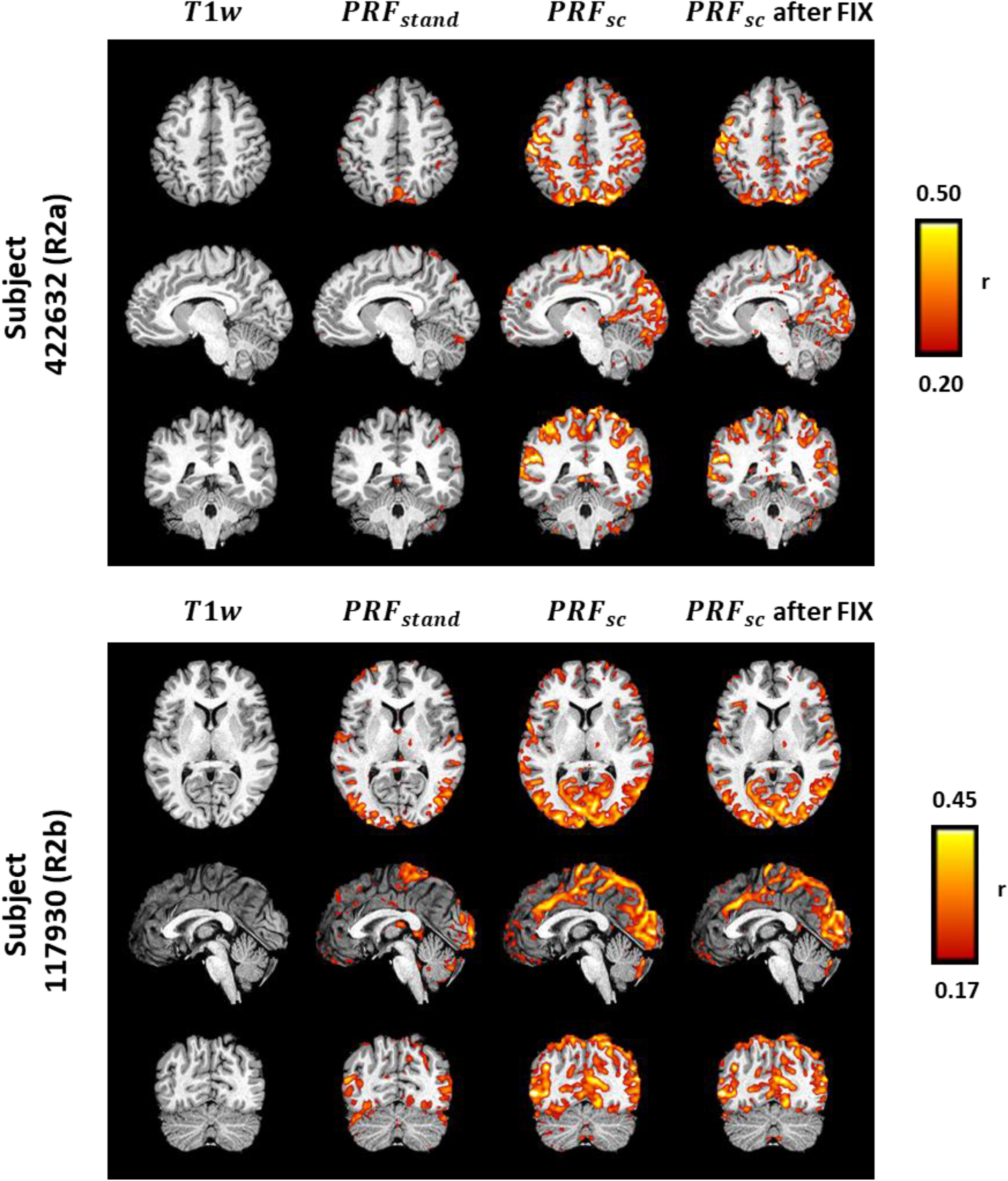
Correlation maps between the physiological model predictions and fMRI timeseries for two representative subjects. (1^st^ column) *T*_1_-weighted images; (2^nd^ & 3^rd^ column) maps derived with the standard (*`PRF`*__*stand*__) and scan-specific (*`PRF`*_*sc*_) model, respectively; (4^th^ column) maps derived with the scan-specific model when applied on data previously corrected with FIX. Overall, all models account for substantial variance in gray matter as well as near large vessels.

The statistical maps shown in Fig. 10 demonstrate much finer detail compared to previous related studies (Birn et al., 2008; Chang et al., 2009; Golestani et al., 2015) which is in part due to the higher spatial resolution (2 mm isotropic voxels) fMRI data acquired in the HCP. However, another factor that may have affected the resolution is the spatial smoothing performed during the preprocessing. To better understand its effect, we extracted these maps without spatial smoothing as well as with a spatial filter with a larger FWHM value (6 mm instead of 3 mm). As shown in Supplementary Figs. 9-10, larger FWHM values resulted in higher correlation values. Overall, our results suggest that a FWHM value of 3 mm yields a good compromise between the resulting signal-to-noise (SNR) ratio and the spatial resolution of the regional maps related to physiological effects on fMRI time-series.

Finally, the contribution of different sources of physiological noise was examined. Specifically, the 6 parameters corresponding to the cardiac-related regressors in RETROICOR (3^rd^ order), estimated during preprocessing, were used to construct the pulsatility-driven component of each voxel time-series, which was subsequently correlated with each voxel time-series to extract the correlation map related to pulsatility. The maps related to HR and RF were extracted separately by employing the scan-specific model (*`CRF`*_*sc*_ and *RRF*_*sc*_ respectively). Fig. 11 shows the contribution of each physiological source for a representative subject on *T*_2_-weighted structural images, instead of the typical *T*_1_-weighted images, as *T*_2_-weighted images yield better contrast for visualizing vessels. We observe that not all areas with large vessels were equally affected by pulsatility or fluctuations in HR and breathing pattern. Furthermore, changes in HR and breathing pattern mostly affected areas in gray matter in the cerebrum, whereas pulsatility affected areas close to the brainstem. Fig. 12 **Fig. 12** illustrates the spatial patterns averaged across all subjects for the three aforementioned physiological noise sources, as well as the spatial patterns for the breathing-related regressors in RETROICOR (3^rd^ order). Note that due to the high collinearity between the set of motion realignment parameters and the set of breathing-related regressors in RETROICOR, the former was orthogonalised with respect to the latter before conducting the GLM, in order to estimate the statistical maps with regions affected by breathing motion. As expected, the breathing-related artifacts were located at the edges of the brain, since the voxels in these regions during breathing move across boundaries of tissue, air or bone that are characterized by large differences in magnetic susceptibility and are, therefore, corrupted by strong motion artifacts. As in Fig. 11, HR and RF effects were found to be more pronounced in distinct areas as compared to cardiac pulsatility. We could not examine whether the changes in HR and breathing patterns were more pronounced in voxels located around draining veins and sinuses rather than arteries, as the HCP does not include images for differentiating veins from arteries. However, a visual comparison with voxel-wise probabilistic maps of veins and arteries developed in (Bernier et al., 2018) suggests that voxels nearby large draining vessels may be mostly affected by changes in HR and breathing pattern, whereas voxels close to arteries may be mostly affected by cardiac pulsatility.

**Fig. 11.**
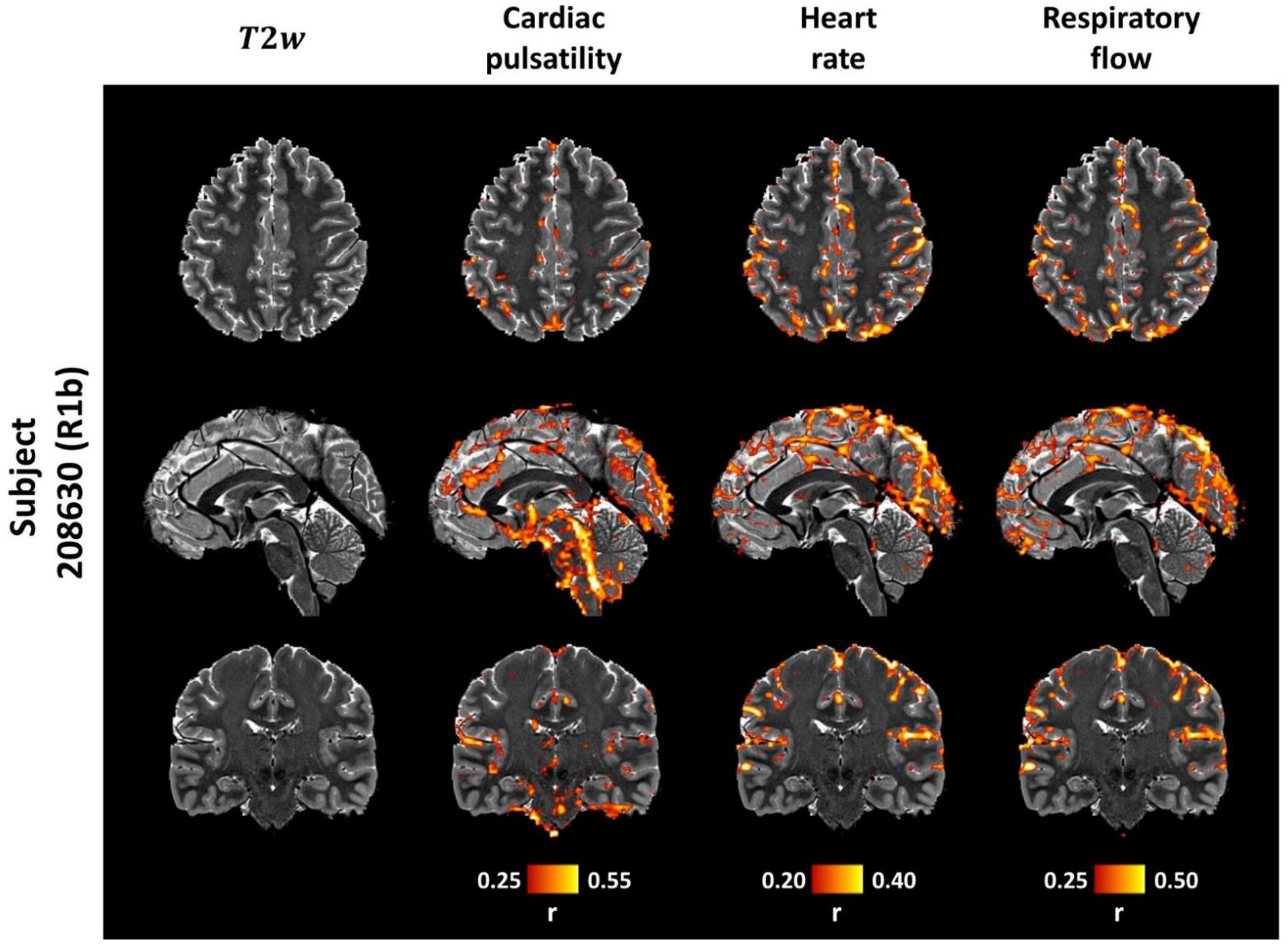
Contribution of different physiological noise sources in fMRI for a representative subject. (1^st^ column) *T*_2_-weighted images; (2^nd^ column) correlation maps related to cardiac pulsatility as modelled with RETROICOR; (3^rd^ column) correlation maps related to HR as modelled with the scan-specific model *`CRF`*_*sc*_; (4^th^ column) correlation maps related to RF as modelled with the scan-specific model *RRF*_*sc*_. While the effect of all physiological sources appears mostly in areas close to vessels, cardiac pulsatility effects are more pronounced around the brainstem, whereas HR and RF effects are more prominent in the occipital and parietal lobes.

**Fig. 12.**
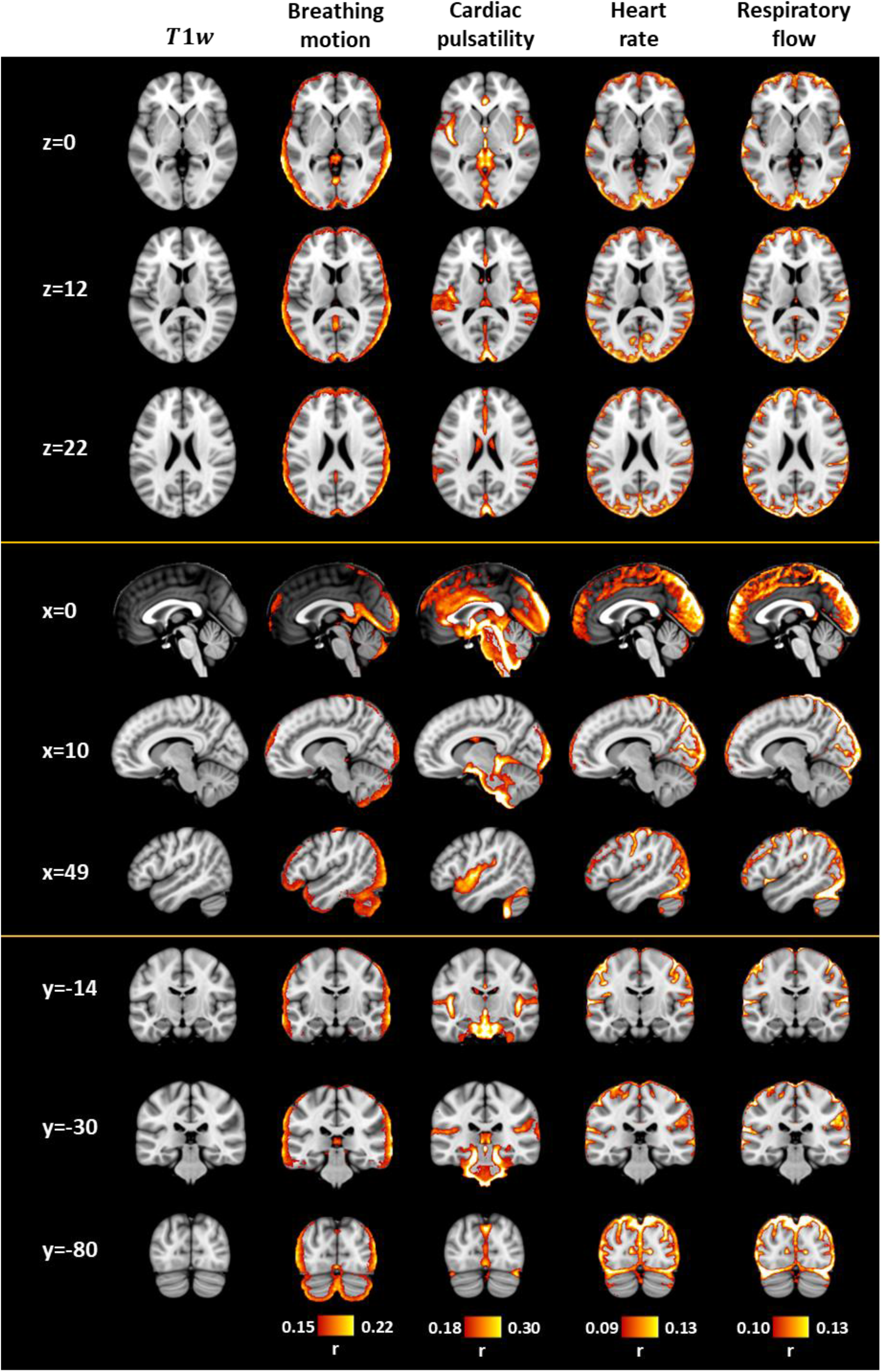
Contribution of different physiological noise sources in fMRI averaged across all subjects. 1^st^ column) MNI152 standard-space *T*_1_-weighted average structural template image (1 mm isometric voxel); (2^nd^ column) correlation maps related to breathing motion as modelled with RETROICOR; (3^rd^ column) correlation maps related to cardiac pulsatility as modelled with RETROICOR; (4^th^ column) correlation maps related to HR as modelled with the scan-specific model *`CRF`*_*sc*_; (5^th^ column) correlation maps related to RF as modelled with the scan-specific model *RRF*_*sc*_. BOLD fluctuations due to breathing motion were found at the edges of the brain as these areas are more prone to susceptibility artifacts. BOLD fluctuations due to cardiac pulsatility were more pronounced close to the basilar and vertebral arteries, in the 4^th^ ventricle, in the superior sagittal sinus, in the lateral sulcus, in the occipital lobe and in the anterior cingulate cortex. On the other hand, BOLD fluctuations due to changes in HR and RF were widespread across gray matter and more pronounced in frontal and posterior brain regions, as well as in sinuses such as the superior sagittal sinus and the transverse sinuses. The correlation maps shown in this figure are available on https://neurovault.org/collections/5654/.

Due to the substantial overlap in the regions affected by HR and RF, we further investigated whether there was a significant correlation between HR and RF, as well as between the corresponding physiological regressors (i.e. HR and RF convolved with *`CRF`*_*sc*_ and *RRF*_*sc*_, respectively) which could explain the spatial overlap. Respiratory sinus arrhythmia (RSA) is the phenomenon where the heart rate is influenced by the breathing cycle, and this could cause HR and RF time-series to be correlated. The peak cross-correlation value between HR and the respiratory signal, which is a measure of RSA, averaged over all scans, was equal to 0.33 ± 0.14 in the data of our cohort. However, the absolute correlation found between HR and RF (i.e. the square of the derivative of the respiratory signal) was lower (0.18 ± 0.10), while the absolute correlation between the corresponding regressors that are considered in the GLM was even lower (0.14 ± 0.10) suggesting that the same brain regions are likely affected by HR and RF. Moreover, we estimated the group-level maps related to HR and RF separately for the scans corresponding to the first quartile of maximum cross-correlation values between HR and the respiratory signal (0.17 ± 0.06) and found again the same regions affected by HR and RF.

### 3.5 Comparison of *PRF* curves for different variants of GS

The results presented above suggest that the scan-specific model *`PRF`*_*sc*_ yields the best performance for modelling the effect of HR and RF. A related question of interest is whether different GS variants significantly affect these results. Specifically, the GS used in the previous sections was defined as the mean time-series from the WB after preprocessing the fMRI data by regressing out the effect of head motion and high-frequency physiological fluctuations (see Methodology). In this section, we considered eight variants of the GS (i.e. WB, GM, WM, CSF, WB_clean_, GM_clean_, WM_clean_ and CSF_clean_) that correspond to the mean time-series from voxels in different ROIs before (no subscript) or after (subscript clean) correcting the fMRI data for head motion and high-frequency physiological fluctuations (i.e., the cardiac-related and breathing-related regressors estimated with RETROICOR). In agreement with previous studies (Power et al., 2017), the WB was almost identical to the GM signal and, based on the variance explained, WB and GM were found to be the most sensitive signals to the effect of HR and RF (Supplementary Fig. 11). Even though the variance explained in the WM and CSF signals was still high (*r* ≈ 0.56), the corresponding *`PRF`*_*sc*_ curves tended to exhibit slower dynamics compared to their GM counterparts (Supplementary Fig. 12). Finally, preprocessing of the fMRI data before deriving the GS signal led to a small decrease in the mean correlation, but it did not affect the shape of the *`PRF`*_*sc*_ curves (Supplementary Figs. 11-12).

### 3.5 Fluctuations in the GS amplitude driven by changes in breathing pattern

In Section 3.3, we associated the variance explained on the GS using the *`PRF`*_*sc*_ models with physiological properties such as HR mean and RF variance within a scan. However, visual inspection revealed that several scans illustrated different breathing patterns even within different time segments of the same scan, suggesting that the variance induced by RF could vary across different segments. While it is beyond the scope of this study to examine time-varying changes in the *`PRF`*_*sc*_ curves or changes in the properties of the GS, exploratory analysis revealed that indeed fluctuations in breathing patterns during the scan were associated to the GS amplitude and GS variance explained by the *RRF*_*sc*_ model. Specifically, as shown in Supplementary Figs.13-14, the more abrupt the breaths were within a specific time segment, the higher the corresponding GS amplitude and the GS variance explained by the *RRF*_*sc*_ model. Furthermore, while in some cases the obtained *RRF*_*sc*_ curves were similar between time segments with different breathing patterns (e.g. scan R1a of subject 201414 shown in Supplementary Fig. 13), in other cases the *RRF*_*sc*_ curves exhibited considerable shape differences across different segments (e.g. scan R2a of subject 207123 shown in Supplementary Fig. 14). However, the differences in *RRF*_*sc*_ curves related to changes in breathing pattern did not seem to follow a specific pattern across scans.

## 4. Discussion

We have rigorously examined the effect of the choice of physiological response functions on the investigation of physiological effects (HR and breathing pattern) on fMRI. To do so, we proposed a novel modeling framework to obtain accurate estimates of the *`PRF`* curves. Linear convolution models, whereby physiological variables are convolved with suitable *`PRF`* curves, were employed to model the associated physiological-driven BOLD fluctuations. The *`PRF`* curves were estimated using numerical optimization techniques that present two main advantages compared to previously used techniques (Birn et al., 2008; Chang et al., 2009; Falahpour et al., 2013; Golestani et al., 2015): 1) they allow the convolution of the physiological variables with the *`PRF`* curves at a high sampling rate independent of the TR of the fMRI data, preventing a possible loss of important information that could happen when a low sampling rate is used. 2) They restrict the shape of the *`PRF`* curves to a specific structure to prevent overfitting, as well as physiologically implausible curves. The structure of the curves was defined as the double gamma function which is also the basis of the canonical HRF in SPM and *RRF*__*stand*__ (Birn et al., 2008), and its parameters were restricted within physiologically plausible ranges. The population-specific models *`PRF`*_*pop*_ demonstrated significantly better fit on the GS as well as in individual voxel time-series compared to the standard models *`PRF`*__*stand*__ (Fig. 6, Fig. 9). The scan-specific model *`PRF`*_*sc*_ outperformed *`PRF`*_*pop*_ while no significant differences were found between the scan-specific model *`PRF`*_*sc*_ and voxel-specific model 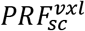. The between-scan variability in the *`PRF`* curves was partly attributed to physiological factors such as the subject’s mean HR during a scan. Overall, HR was found to explain higher frequency fluctuations on the GS than RF (Fig. 4). Consistent with previous findings, changes in HR and breathing pattern (breathing rate and depth) had a strong effect across widespread regions in the gray matter (Birn et al., 2008; Chang et al., 2009).

In practice, the scan specific model *`PRF`*_*sc*_ that demonstrated the best performance, can be easily applied to fMRI data as long as concurrent physiological recordings are provided and the examined scan lasts more than 5 minutes. However, if the duration of the data is short and there is the risk of overfitting with *`PRF`*_*sc*_ (Fig. 7b), the population-specific model *`PRF`*_*pop*_ should be preferred that accounts for potential differences in the curves compared to the ones obtained in the present study related to the population examined or due to a different scanner and pulse sequence. The Matlab codes for the *`PRF`*_*sc*_ model as well as the rest of the models examined here can be found on https://github.com/mkassinopoulos/PRF_estimation. The *`PRF`* parameters can be estimated from the mean time-series of voxels in WB, GM, WM or CSF. However, the GM and WB mean time-series, which are almost identical and are typically used to define the GS, were found to be more strongly correlated to HR and RF fluctuations, compared to the WM and CSF time-series (Supplementary Fig. 11). Using the WM and CSF mean time-series to define the GS also yielded *`PRF`*_*sc*_ curves with slower dynamics (Supplementary Fig. 12). Therefore, in the case that the signal of interest is in GM regions, using the GM or WB compartment to extract the GS is a more suitable choice for *`PRF`* parameter estimation. The GS needed for the parameter estimation, can be obtained directly from the fMRI data, after the basic preprocessing of volume realignment and high-pass filtering.

### 4.1 Population specific vs standard *PRF* curves

Using the proposed framework, we derived population specific curves (*`CRF`*_*pop*_ and *RRF*_*pop*_), which explained a substantially larger fraction of GS and individual voxel time-series variance compared to the standard models *`PRF`*__*stand*__ (Fig. 6, Fig. 9). The population-specific HR model *`CRF`*_*pop*_ demonstrated considerably faster dynamics than *`CRF`*__*stand*__ (Chang et al., (2009); Fig. 2a). Some of the main differences in the two studies that may explain the results are the following: a) in Chang et al., (2009) the resting-state scan was performed with eyes closed instead of eyes open as done in the HCP data used here, and b) *`CRF`*__*stand*__ was obtained by first estimating the *`CRF`* with the method of maximum a posteriori in individual voxels and then averaging across voxels and subjects, whereas, in the present study, similar to Falahpour et al. (2013), *`CRF`* estimation was based on the GS of each scan, which is partly driven by physiological noise, and, thus, yields an adequate SNR. In addition to these, one of the reasons for the faster dynamics observed in the `CRF` curves in the present study is the fact that in previous studies (Chang et al., 2009; Falahpour et al., 2013; Golestani et al., 2015) HR was initially averaged within a time window of 4-6 s and subsequently downsampled to a low TR (e.g. 3 s) before further analysis, disregarding fast fluctuations in HR. In contrast, the framework employed here has the advantage that it can perform the convolution of the HR with the *`PRF`* curve at a higher sampling rate and downsampling occurs at the output stage for comparison with the measured fMRI time-series (i.e. the GS in our case).

Differences, albeit to a smaller extent, were also found between the population-specific curve *RRF*_*pop*_ and the standard curve *RRF*__*stand*__ (Birn et al., 2008; Fig. 3d). However, in the proposed model, a different feature (RF) was used as an input and, thus, direct comparisons between *RRF*_*pop*_ and *RRF*__*stand*__ should be made with caution. The RF was introduced here as a physiological variable derived from the respiratory signal and was preferred to RVT (Birn et al., 2008) as it does not require peak detection, a task that is not always straightforward for respiratory signals as the breathing volume and rate during spontaneous breathing can vary over a wide range across time. A possible explanation for the increased performance of the proposed *RRF*_*pop*_ model compared to the standard method is that *RRF*_*pop*_ was estimated using resting-state data with the RF and the GS as the input and output of the model, whereas in Birn et al. (2008), *RRF*__*stand*__ was derived from the BOLD response induced by a deep breath without incorporating the RVT in the estimation stage. Note that Birn et al. (2008) also reported a poor fit of their method in resting-state fMRI, which was improved only when allowing time-shifting separately for each voxel, an approach that was followed later by (Bianciardi et al., 2009; Chang and Glover, 2009b). Time shifting is a reasonable choice in physiological modelling as it can take into consideration time lags between measurements obtained from different instruments (e.g. fMRI time-series from the MRI scanner vs end-tidal gas measurements from a gas analyzer with a long sampling line), as well as differences in the blood time arrival between distant brain regions due to differences in vascular pathways (Bright et al., 2009). However, time shifting has been shown to inflate the correlation statistics and, therefore, validation of the optimal temporal shift is needed in future studies (Bright et al., 2016).

The population specific curve *`CRF`*_*pop*_ (Fig. 3a) was characterized by a peak at 1.2 s and an undershoot at 7.0 s. A possible explanation about the first peak is that increases (decreases) in HR are briefly followed by an increase (decrease) in cardiac output and, in turn, in CBF and BOLD signal. On the other hand, the undershoot may indicate the presence of a negative feedback mechanism (e.g. decrease of stroke volume and, thus, cardiac output or decrease in relative distribution of cardiac output to the cerebral vasculature) which ensures that CBF is maintained at normal levels. A positive peak followed by a negative peak was observed for *RRF*_*pop*_ as well (Fig. 3d). A possible explanation for this behavior is the following: Increases in RF are followed by increases in the levels of O2 in cerebral blood which in turn lead to a decrease in levels of deoxygenated blood and an increase in the BOLD signal. However, increases in RF are also followed by decreases in levels of CO_2_ in the blood, which is a strong vasodilator. As a result, decreases in levels of CO_2_ are followed by a decrease in CBF and BOLD signal. This vasoconstriction, however, is likely a slower process, which can explain the decrease in BOLD signal with a minimum peak at about 13 seconds after the RF increase. In a similar manner, a decrease in RF would lead to an initial decline in BOLD signal followed by a slow overshoot.

### 4.2 Variability in *PRF* curves across subjects and scans

Among the different physiological models examined in this study, the scan specific model *`PRF`*_*sc*_ yielded the best performance (Fig. 6, Fig. 9). This suggests that physiological response functions do not only vary across subjects but also across scans of the same subject. Importantly, our results suggest that scan-specific curves can be robustly derived from whole-brain resting-state fMRI data with durations longer than 5 minutes and high sampling rate (e.g. TR=0.72) (Fig. 7b). As the physiological origin of the *`CRF`* and *RRF* curves is different, their form is discussed separately.

Visual inspection of the scan-specific curves *`CRF`*_*sc*_ revealed differences across subjects and across sessions within-subjects. A more systematic comparison revealed that the time of the negative peak of the curve was strongly dependent on the mean HR of the subject, with shorter times linked to higher mean HR (Fig. 5a). As the mean HR was found to vary significantly between the two sessions within-subject (Fig. 2), this may explain the differences in *`CRF`*_*sc*_ curves across sessions. Differences in *`CRF`*_*sc*_ curves across sessions within-subject could be attributed also to differences in arterial blood pressure that may vary significantly between scans collected at different days. However, we could not examine this factor as blood pressure measurements were not collected at the day of each fMRI scan. The finding that the time of negative peak of the curve was associated with higher mean HR may suggest that when HR is higher, CBF is also higher and a fast negative feedback is more critical to prevent extreme values of CBF.

In our work, we assumed that the relationship between HR and the BOLD signal can be described with a linear time-invariant system. However, it may be the case that a time-varying system, the parameters of which are expressed as a function of the time-varying mean (or instantaneous) HR and possibly blood pressure may explain the fluctuations in GS more accurately. Moreover, we observed that *`CRF`*_*sc*_ explained a larger fraction of GS variance for scans with a low mean HR. Subsequent analysis showed a strong positive correlation between the mean HR and fluctuations in HR (results not shown here), which could indicate that a time-varying *`CRF`* is more appropriate for cases where HR is high and varies significantly. On the other hand, a possible explanation that scans with high mean HR yielded a weaker correlation between fluctuations in HR and GS is that in these scans the subjects were more stressed, which explains the high mean HR, and tended to move more, distorting the fMRI data (including the GS). To confirm whether head motion was linked to the mean HR, we calculated the framewise displacement (FD), which reflects the relative head motion during the scan (Power et al., 2012) using the motion realignment parameters. The mean FD was found to be strongly correlated to the mean HR (*r* = 0.20; *p* = 0.007) supporting the above interpretation. Additionally, another explanation for the negative correlation between mean HR and GS variance explained by the HR is that subjects with high mean HR were in a state of higher arousal levels and, consequently a significant component of the GS fluctuations was neuronally-driven. However, we could not examine the validity of this interpretation with our data as we did not have concurrent data (e.g. EEG data) that can provide an index of arousal levels.

The population specific curve *RRF*_*pop*_ demonstrated relatively mild differences compared to the *RRF*__*stand*__. On the other hand, the scan specific curves *RRF*_*sc*_ exhibited a larger degree of variability across subjects and scans compared to *`CRF`*_*sc*_, with curves for a large number of scans having only negative values, instead of the commonly observed positive peak followed by an undershoot (Birn et al., 2008; Chang et al., 2009; Power et al., 2017). These differences could not be attributed to the different individual patterns in BR or RF. Nonetheless, the proportion of GS variance explained with RF was found to be correlated with RF variance (Fig. 5c). RF was defined as the squared derivative of the respiratory signal, which has a similar form with the FD measures proposed in the literature for a surrogate of relative head motion (Power et al., 2012; van Dijk et al., 2012). Head motion is a main source of noise in fMRI, as it can cause spin history related motion artifacts in the BOLD signal with finite time memory (Friston et al., 1996). In this context, RF could be viewed as an index of relative head motion induced by breathing that is sampled at a higher sampling rate compared to the motion parameters estimated in fMRI preprocessing during volume realignment (see Supplementary Fig. 14 for an example of a scan with high similarity between RF and FD waveforms). Interestingly, a recent paper used the FD to examine temporally-lagged artifacts on the GS due to head motion (Byrge and Kennedy, 2018) and reported impulse responses describing the effects of the latter similar to the *RRF*_*pop*_ curve in the present paper. They also found, based on concurrent respiratory signals, that the FD fluctuations were strongly associated to breathing. Therefore, apart from fluctuations due to changes in CO_2_ levels, RF, through convolution with the *RRF*_*sc*_, can potentially remove residuals of breathing-related motion artifacts that cannot be removed completely with the motion realignment parameters or RETROICOR regressors through linear regression. The aforementioned motion artifacts are caused by head movement and variations in the air volume inside the lungs when the subject breathes, leading to fluctuations in the static magnetic field that are ultimately reflected on the fMRI data. Moreover, the breathing-related motion artifacts may be related to the body type and breathing behavior of each subject, as well as the position of the subject inside the MRI tube, which could explain the variability of the *RRF*_*sc*_ curves across subjects and sessions within-subject and the improved performance of the scan-specific model compared to the population-specific model *RRF*_*pop*_, particularly in scans with high RF variance.

Power et al. (2017) have previously shown that the standard deviation of the GS is positively correlated to the variability in HR and respiration as well as to the extent of motion during the scan. In this study, we were able to replicate these findings (results not shown) but also show evidence that changes in the breathing pattern across the scan can alter the shape in the *RRF* curves as well as modulate the amplitude of the global signal (Supplementary Figs. 13-14). This finding suggests that the time-varying changes in the GS amplitude driven by fluctuations in cardiac and breathing activity may play an important role in the apparent time-varying rs-connectivity (Preti et al., 2017) and, thus, should be taken into consideration in these studies.

### 4.3 Physiological noise correction with FIX and GSR

FIX is a widely-used tool for denoising fMRI data, as it removes fluctuations due to motion and cardiac pulsatility in an automated manner without the need for physiological recordings (Salimi-Khorshidi et al., 2014). HCP has been using it to provide FIX-denoised data and many researchers have been analyzing these data without any further preprocessing (Bijsterbosch et al., 2017; Vidaurre et al., 2017). However, Burgess et al. (2016) have recently demonstrated, using grayordinate plots, that while FIX-denoising substantially reduces spatially specific artifacts, it yields only a mild decrease in global fluctuations.

Our study provides further evidence that FIX does not completely remove global artifacts, and particularly fluctuations due to changes in HR and breathing pattern. The effects of these fluctuations are typically widespread within the gray matter and mainly in frontal and posterior brain regions (Fig. 12). As a side note, our analysis showed that breathing motion and cardiac pulsatility artifacts that are often modelled with RETROICOR are efficiently removed with FIX (see Supplementary Fig. 15 for correlation maps from representative scans related to RETROICOR regressors before and after using FIX). It is well established that spontaneous fluctuations in physiological processes lead to global fluctuations in the fMRI BOLD signal (Chang et al., 2009). Furthermore, these fluctuations may also have a significant effect in rs-FC analyses, including dynamic rs-connectivity (Birn, 2012; Murphy et al., 2013; Nikolaou et al., 2016) and, therefore, should be taken into consideration in the analysis. However, it has been suggested that spatial ICA, which is used in FIX as well as in other ICA-based denoising techniques such as ICA-AROMA (Pruim et al., 2015) is mathematically, by design, unable to separate global temporal artifacts from fMRI data (Glasser et al., 2018).

The GS, which is simply the BOLD time-series averaged across all voxels in the brain or gray matter, is often regressed out from the fMRI data, in conjunction with other nuisance regressors (Aguirre et al., 1997; Fox et al., 2005) or FIX (Burgess et al., 2016; Siegel et al., 2017), in order to correct for global artifacts. GSR has been shown to improve the correspondence of properties of fMRI rs-FC with observations from neuroanatomy (Fox et al., 2009) and to substantially reduce motion-group differences (Burgess et al., 2016). However, there is no consensus yet in the field whether GSR should be performed (Liu et al., 2017; Murphy and Fox, 2017). Murphy et al. (2009) was the first study to mathematically demonstrate that GSR introduces spurious anticorrelations in rs-FC. Moreover, many recent studies have reported strong correlations between the fluctuations or amplitude of GS and neuronal-related measures such as electrophysiological activity from intracranial recordings (Scholvinck et al., 2010) and vigilance levels (Chang et al., 2016; Falahpour et al., 2018; Wong et al., 2016, 2013). As a result, to facilitate interpretation, reviewers have been recommending the repetition of rs-FC studies with and without GSR to address whether the results can be attributed to GSR or not, while a recent study has developed a measure termed Impact of the Global Average on Functional Connectivity (IGAFC), which quantifies the extent of the impact of GSR on inferences based on seed-based statistical correlation maps (Carbonell et al., 2014). Furthermore, researchers have proposed alternatives to GSR. For example, Glasser et al. (2018) have recently proposed the use of temporal ICA after FIX denoising to preserve the neuronal-related component of the global signal, while removing global structured noise. However, this technique is only applicable to large datasets such as HCP. In addition, Carbonell et al. (2011) have proposed a method based on PCA for regressing out global artifacts and fluctuations that are uncorrelated to network-specific activity, even though it cannot ensure the preservation of global neurophysiological activity.

Our results suggest that a good alternative to GSR are the *`PRF`* models proposed here. As the physiological regressors are extracted from concurrent physiological recordings, in contrast to GSR and other data-driven techniques, they can account for fluctuations that are mainly physiologically-driven. However, resting physiological fluctuations may influence neuronal activity; for instance, a recent magnetoencephalography (MEG) study has shown that spontaneous changes in arterial CO_2_ have an influence on neural activity (Driver et al., 2016). Therefore, the risk of removing some neuronal-related fluctuations cannot be completely excluded. The *`PRF`*_*sc*_ models can be trained on a scan-by-scan basis based on the GS and physiological recordings of cardiac activity and breathing. The parameter estimation for all 164 scans examined here required in total about 3 minutes on a desktop computer with Intel® Core™ i7-5820K (3.3 GHz) CPU and 64 GB RAM. The extraction of the GS can be done by averaging the time-series from voxels either within the WB or GM of the raw fMRI data (i.e. after the basic preprocessing of volume realignment and high-pass filtering). Following the training, the physiological regressors are extracted and can be subsequently removed from the data through linear regression or added in the general linear model as regressors along with other regressors of interest. Importantly, the framework proposed in this study for estimating the *`PRF`* curves performs the convolution of physiological variables with the *`PRF`* curves at a higher sampling rate than the TR of the fMRI acquisition and, therefore, can model the high-frequency effects of HR even with a long TR (~3-4 s). The codes for the *`PRF`* models presented here are publicly available and can be found on https://github.com/mkassinopoulos/PRF_estimation whereas the group-level correlation maps related to HR, RF, cardiac pulsatility and breathing motion artifacts (Fig. 12) are available on https://neurovault.org/collections/5654/.

In future work, we aim to extend the proposed framework in the context of task-based fMRI, in which physiological processes are often modulated by the experimental paradigm (Glasser et al., 2018) and, thus, physiologically-driven fluctuations may be interpreted as neuronally-driven fluctuations and vice versa. If the *`PRF`*_*sc*_ curves are estimated from the GS of the task-based scan, without incorporating the information about the experimental paradigm, the *`PRF`*_*sc*_ curves may be biased by the HRF and, subsequently, physiological confounds as well as signal of interest from the fMRI data may be removed. On the other hand, if the *`PRF`*_*sc*_ curves are estimated from a separate resting-state scan, changes in the *`PRF`* curves associated with a different physiological state (e.g. higher mean HR) during the task compared to the resting-state scan will not be taken into consideration in the model and, therefore, physiologically-driven fluctuations will not be well modelled.

## 5. Conclusion

In this study, we have developed a novel method for removing the effect of fluctuations in HR and breathing pattern in BOLD fMRI data by combining optimization and basis expansion techniques for the robust estimation of subject and scan-specific *`PRF`*_*sc*_ curves. Our approach was validated on data from the Human Connectome Project (HCP) and achieved improved performance compared to current methods, including the standard *`CRF`*__*stand*__ and *RRF*__*stand*__, and FIX.

The proposed framework is of great interest for researchers interested in studying rs-FC in groups where breathing, heart rhythms or cerebrovascular reactivity can differ. Ultimately, better understanding and quantifying physiological effects on fMRI studies can pave the way for understanding the normal and pathological brain as well as accelerate the discovery of connectivity-based biomarkers for diagnosing neurological disorders, as it will contribute towards disentangling the neural vs. physiological sources of rs-FC.

## Acknowledgments

This work was supported by the Natural Sciences and Engineering Research Council of Canada (Discovery Grant 34362 awarded to GDM), the Fonds de la Recherche du Quebec - Nature et Technologies (FRQNT; Team Grant PR191780-2016 awarded to GDM) and the Canada First Research Excellence Fund (awarded to McGill University for the Healthy Brains for Healthy Lives initiative). MK acknowledges funding from Québec Bio-imaging Network (QBIN). Data were provided by the Human Connectome Project, WU-Minn Consortium (Principal Investigators: David Van Essen and Kamil Ugurbil; 1U54MH091657) funded by the 16 NIH Institutes and Centers that support the NIH Blueprint for Neuroscience Research; and by the McDonnell Center for Systems Neuroscience at Washington University.

## Footnotes

1 Note that the term respiration in physiology refers to the exchange of oxygen and CO_2_ across a membrane, either in the lungs or at the cellular level. However, to be consistent with the terms used in the fMRI literature, throughout the text, by respiration we refer to the act of breathing or ventilation, which is defined as the process of moving air into and out of the lungs.

