## Supplementary Material for "Identification of Physiological Response Functions to Correct for Fluctuations in Resting-State fMRI related to Heart Rate and Respiration"

**Supplementary Table 1.** Subjects included in the study

| Subject | Sex | Age | Subject | Sex | Age |
|---|---|---|---|---|---|
| 100307 | Female | 26-30 | 203923 | Female | 26-30 |
| 105115 | Male | 31-35 | 204622 | Female | 26-30 |
| 108121 | Female | 26-30 | 207123 | Male | 26-30 |
| 113922 | Male | 31-35 | 207426 | Male | 26-30 |
| 117930 | Female | 31-35 | 208630 | Male | 31-35 |
| 118528 | Female | 26-30 | 209127 | Female | 31-35 |
| 118730 | Male | 22-25 | 210011 | Male | 22-25 |
| 118932 | Male | 26-30 | 210415 | Female | 26-30 |
| 120212 | Female | 31-35 | 210617 | Female | 31-35 |
| 120717 | Female | 31-35 | 300719 | Male | 26-30 |
| 121416 | Male | 26-30 | 304020 | Male | 31-35 |
| 121618 | Male | 31-35 | 305830 | Female | 22-25 |
| 121921 | Male | 31-35 | 307127 | Female | 31-35 |
| 122620 | Male | 26-30 | 309636 | Female | 26-30 |
| 122822 | Female | 31-35 | 406432 | Male | 26-30 |
| 200008 | Female | 26-30 | 412528 | Female | 31-35 |
| 200614 | Female | 31-35 | 421226 | Male | 31-35 |
| 200917 | Female | 22-25 | 422632 | Female | 22-25 |
| 201414 | Female | 22-25 | 424939 | Male | 26-30 |
| 201818 | Female | 26-30 | 555348 | Female | 31-35 |
| 203418 | Male | 26-30 | | | |

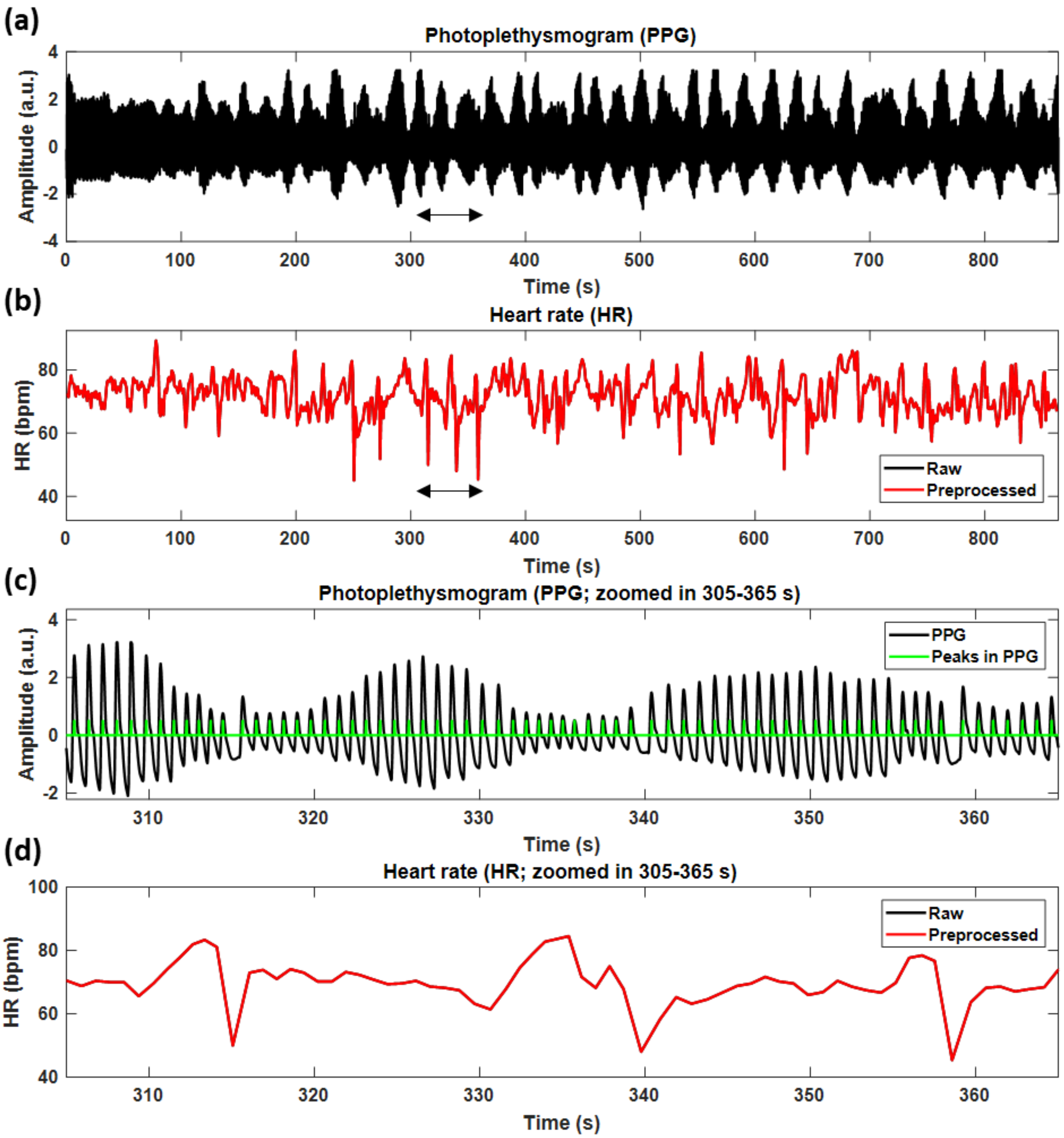


**Supplementary Fig. 1.** Example of heart rate (HR) signal that was not corrected for outliers (scan S207123-R1a). (a) Photoplethysmogram (PPG) used to detect heartbeats and yield HR signal. (b) Raw (black) and preprocessed (red) HR signals shown for the entire scan. (c) PPG zoomed in on a time segment with abrupt changes in HR. (d) HR zoomed in on the same time segment with (c). For this scan, even though there were abrupt changes in HR, the raw HR was used in the analysis without any further preprocessing as, based on visual inspection, the peaks in the PPG were correctly detected by the algorithm.

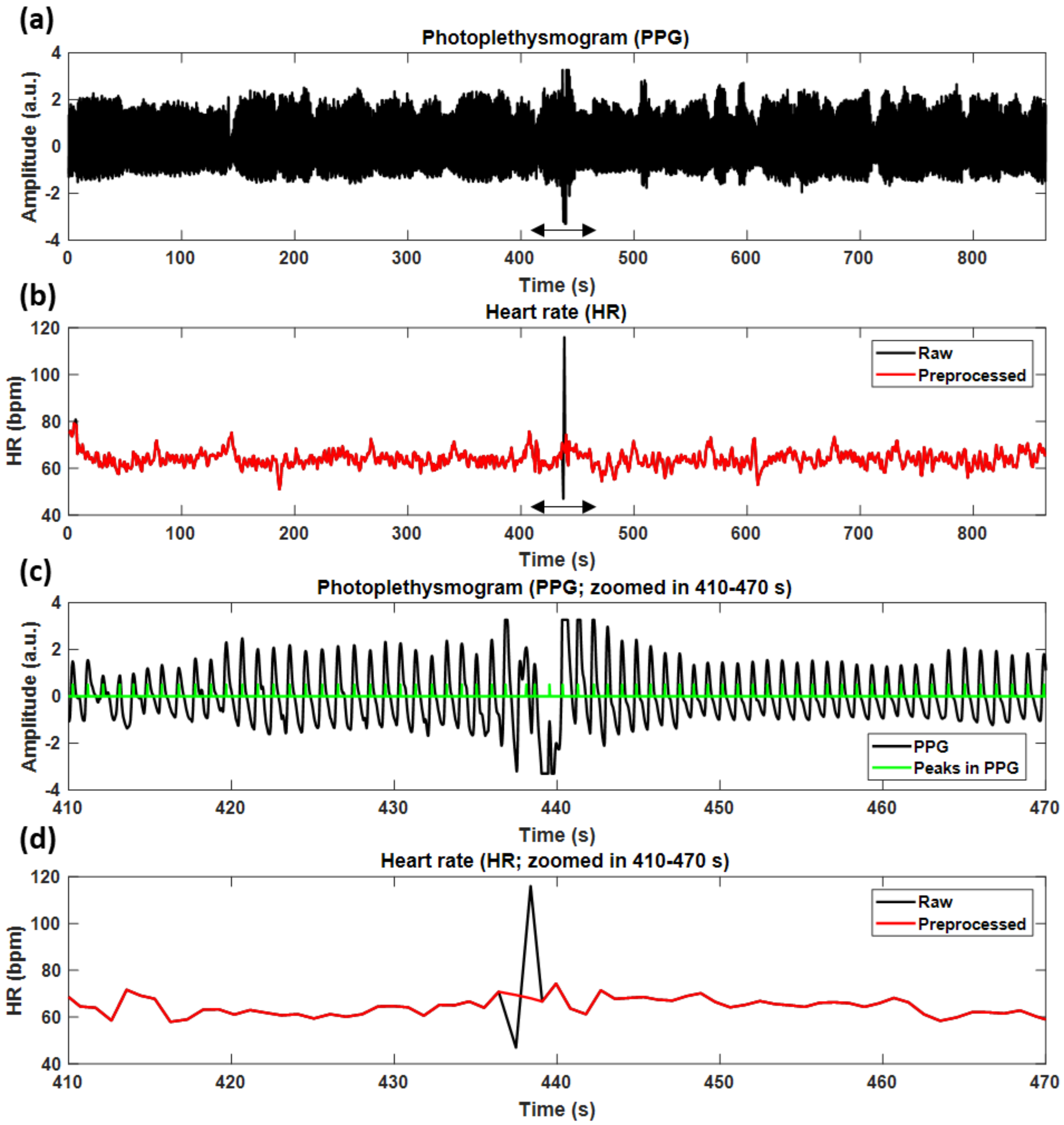


**Supplementary Fig. 2.** Example of heart rate (HR) signal that was corrected for outliers (scan S118730-R2b). (a) Photoplethysmogram (PPG) used to detect heartbeats and yield HR signal. (b) Raw (black) and preprocessed (red) HR signals shown for the entire scan. (c) PPG zoomed in on a time segment with abrupt changes in HR. (d) HR zoomed in on the same time segment with (c). We observe that there are abrupt changes in HR at around 438 s that are due to a noisy segment in the PPG. Therefore, for this scan, the outliers in the raw HR signal were detected as described in section 2.2.1 and replaced using linear interpolation (for this scan, outliers were defined as the time points that deviated more than 5 median absolute deviations from the moving median value within a time window of 30 seconds).

**Supplementary Table 2.** Pseudocode of the algorithm used for $PRF$ parameter estimation (obj_function: objective function, $PRF$ parameters: G=[t1c,d1c,t2c,r2c] and R=[Rc,Rr], HR: heart rate, RF: respiratory flow, GS: global signal, conv: convolution, const: constant term, lin: linear term, corr: correlation coefficient)

| **Pseudocode for models $PRF_{pop}$ to $PRF_{sbj,sdd}$ (Table 1)** | **Pseudocode for models $PRF_{pop}^{sc}$ to $PRF_{sc}$ (Table 1)** |
|---|---|
| function [obj_function] = optPRF_GRpar(G,R, [HR, RF & …<br>GS] of all related scans according to the examined model)<br><br>CRF = gamma(t1c,d1c) + Rc*gamma(t2c,d2c)<br>RRF = gamma(t1r,d1r) + Rr*gamma(t2r,d2r)<br><br>for scan = 1 to N_scans<br>y = GS(scan)<br><br>xCard = conv(CRF, HR(scan))<br>xResp = conv(RRF, RF(scan))<br>X = [const, lin, xCard, xResp]<br><br>X = downsample_to_fMRI_FS(X)<br>B = X\y<br>yPred = xRegr*B<br>r(scan)= corr(y,yPred)<br>end<br><br>obj_function = mean(r) | function [obj_function] = optPRF_Gpar(G, [HR, RF & …<br>GS] of all related scans according to the examined model)<br><br>CRF1 = gamma(t1c,d1c)<br>CRF2 = gamma(t2c,d2c)<br>RRF1 = gamma(t1r,d1r)<br>RRF2 = gamma(t2r,d2r)<br><br>for scan = 1 to N_scans<br>y = GS(scan)<br><br>xCard1 = conv(CRF1, HR(scan))<br>xCard2 = conv(CRF2, HR(scan))<br>xResp1 = conv(RRF1, RF(scan))<br>xResp2 = conv(RRF2, RF(scan))<br>X = [const, lin, xCard1, xCard2, xResp1, xResp2]<br><br>X = downsample_to_fMRI_FS(X)<br>B = X\y<br>yPred = xRegr*B<br>r(scan)= corr(y,yPred)<br>end<br><br>obj_function = mean(r) |

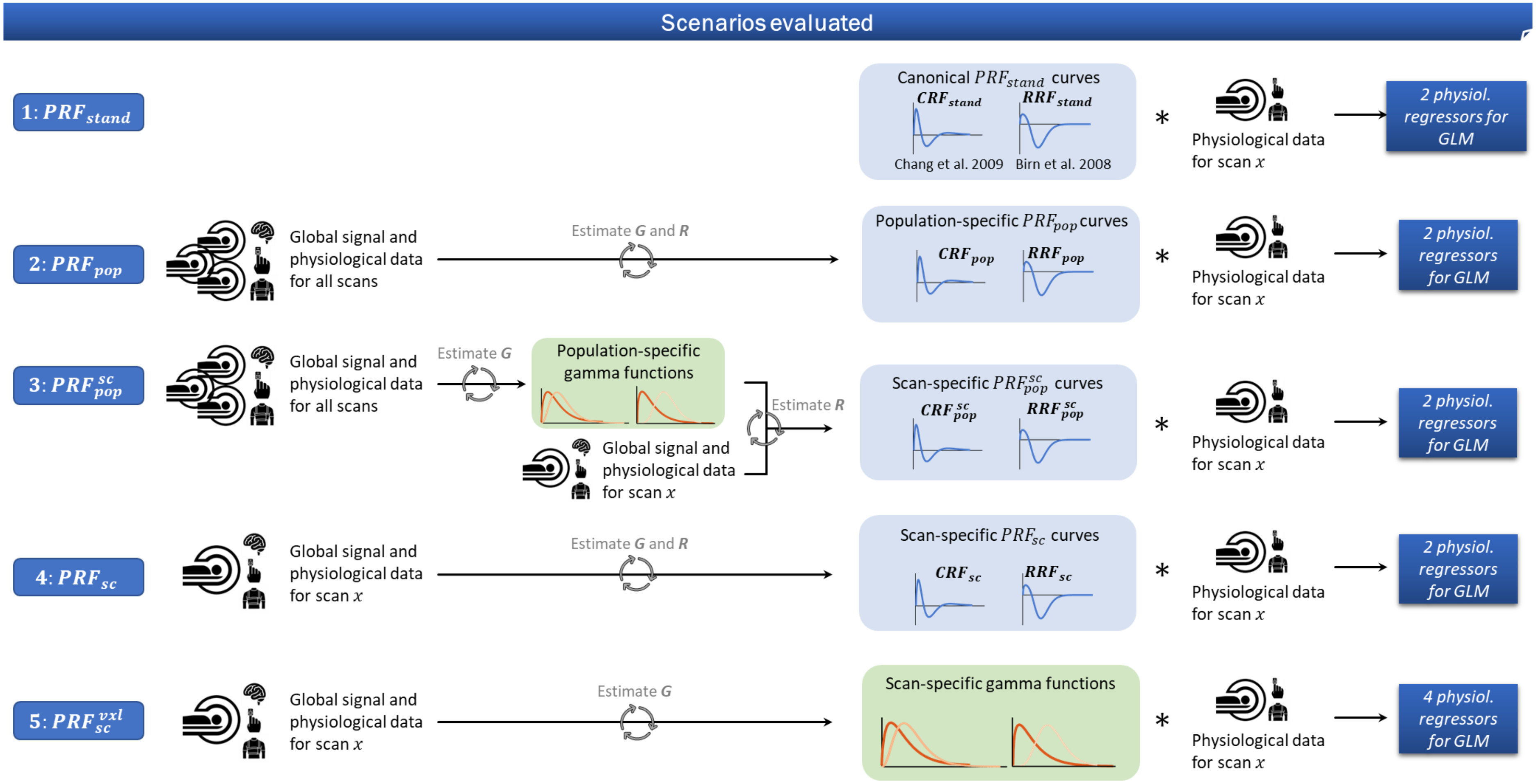


**Supplementary Fig. 3.** Illustration of the required steps for deriving the physiological regressors for the voxel-wise GLM analysis in the five main examined models. The cross-validation framework is omitted for simplicity. Also, the models of the form $PRF_{sbj,xxx}$ and $PRF_{sbj,xxx}^{sc}$ from groups II and III in Table 1 respectively, are omitted from this diagram as they are variations of the models $PRF_{pop}$ and $PRF_{pop}^{sc}$ from the groups II and II, respectively (Table 1). Specifically, they only differ in the fact that the data considered in the first step for the estimation of the curves and gamma functions are obtained from the same subject rather than the same population. The codes for employing all these five models can be found on https://github.com/mkassinopoulos/PRF_estimation.

**Supplementary Figs. 4-7: Demonstration of the $PRF_{sc}$ model for all four scans of two subjects (210415 and 307127)**

Steps: First, the heart rate (HR; 1st row of each scan's panel) and respiratory flow (RF; 5th row) were extracted from the physiological signals. Subsequently, $\boldsymbol{G}$ and $\boldsymbol{B}$ that define the subject-specific $CRF_{sc}$ and $RRF_{sc}$ curves were estimated. The physiological regressors $X_{HR}$ and $X_{RF}$ shown in the 2nd and 4th row were modelled as the convolution of the HR and RF with the $CRF_{sc}$ and $RRF_{sc}$ curves, respectively. Last, the linear combination of the physiological regressors given by $\boldsymbol{B}$ was used to maximize the explained variance on the GS.

In general, we observed that the HR explained higher frequency fluctuations on the GS than RF while their contribution on explained variance varied across scans and subjects. For example, RF tended to explain more variance than HR for subject 210415 whereas the opposite trend appeared for subject 307127. Moreover, we noticed significantly different $RRF_{sc}$ curves between the two subjects. The figures of the scan-specific models $PRF_{sc}$ for all scans are available for inspection in our figshare repository (https://doi.org/https://doi.org/10.6084/m9.figshare.c.4585223.v4; Kassinopoulos and Mitsis, 2019).

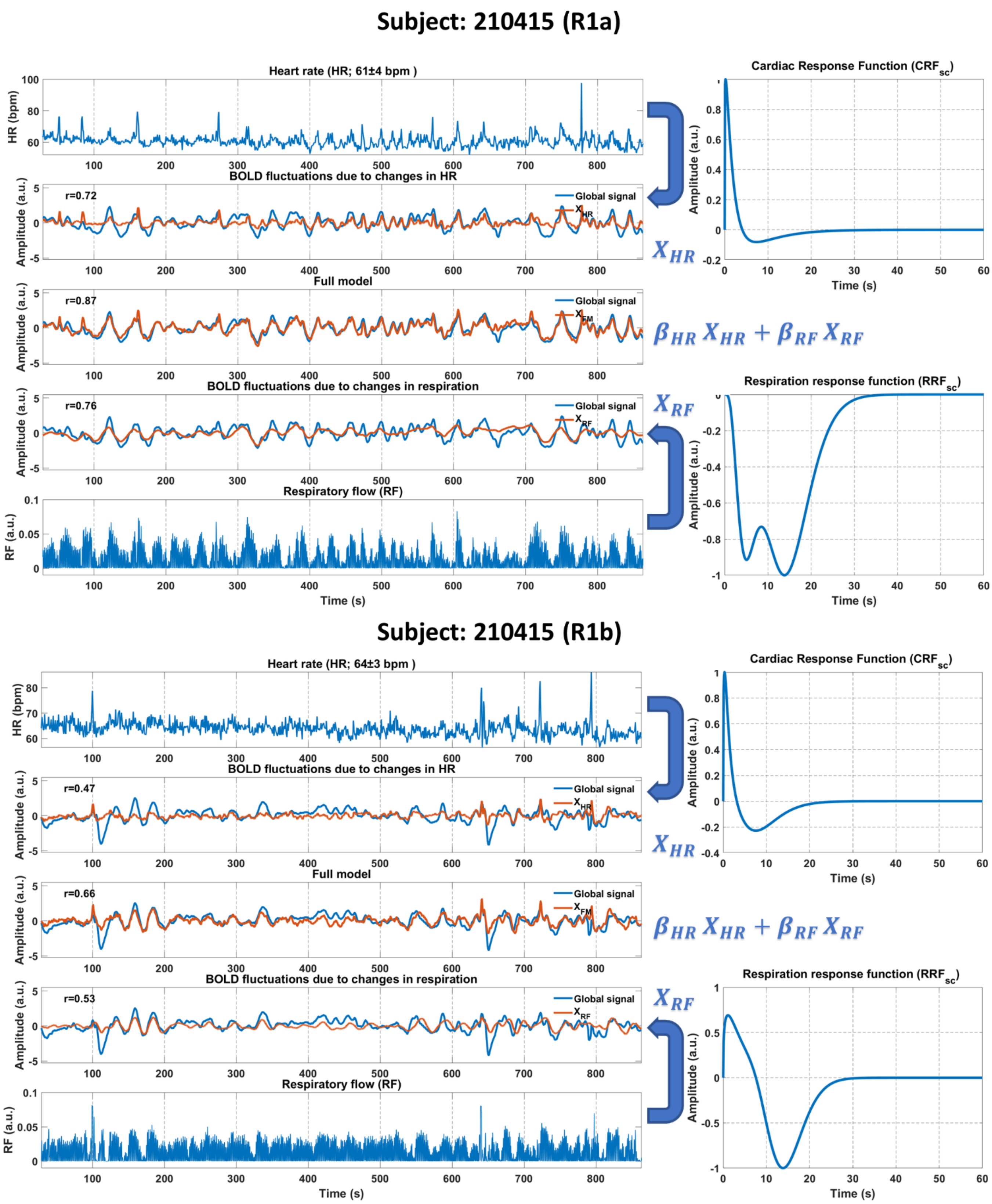


**Supplementary Fig. 4.** Demonstration of the $PRF_{sc}$ model for scans from the 1st session of subject 210415

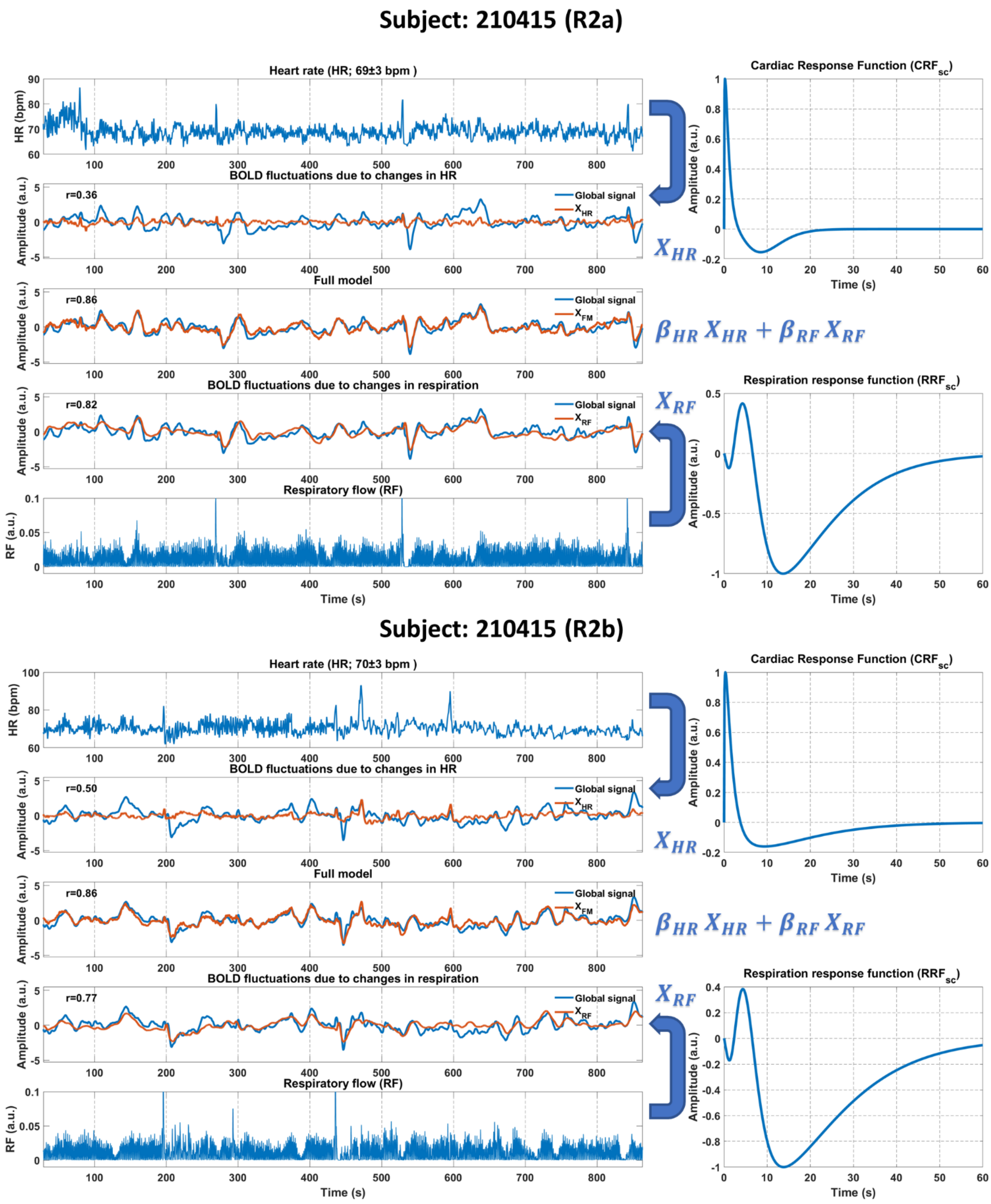


**Supplementary Fig. 5.** Demonstration of the $PRF_{sc}$ model for scans from the 2nd session of subject 210415

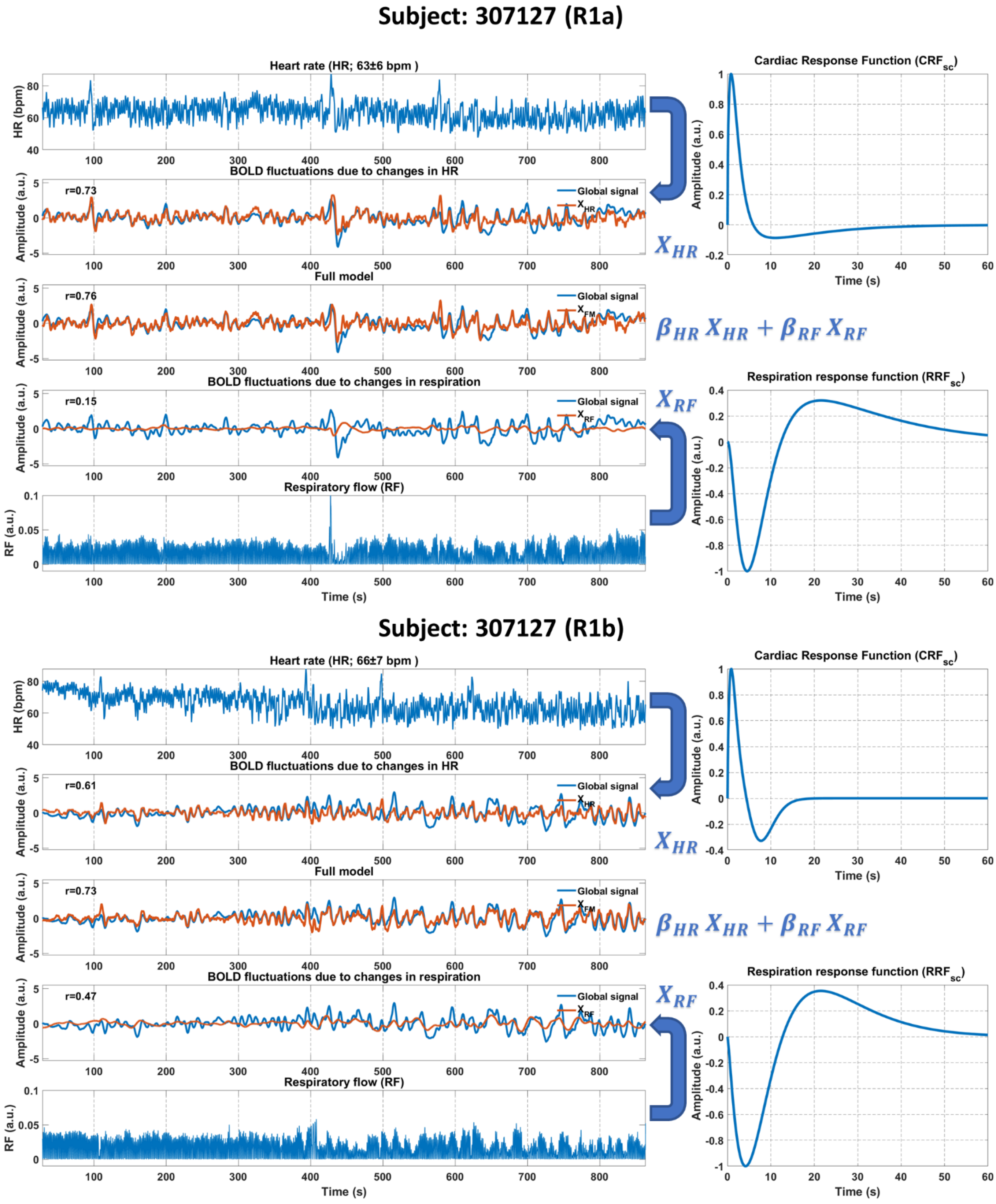


**Supplementary Fig. 6.** Demonstration of the $PRF_{sc}$ model for scans from the 1st session of subject 307127

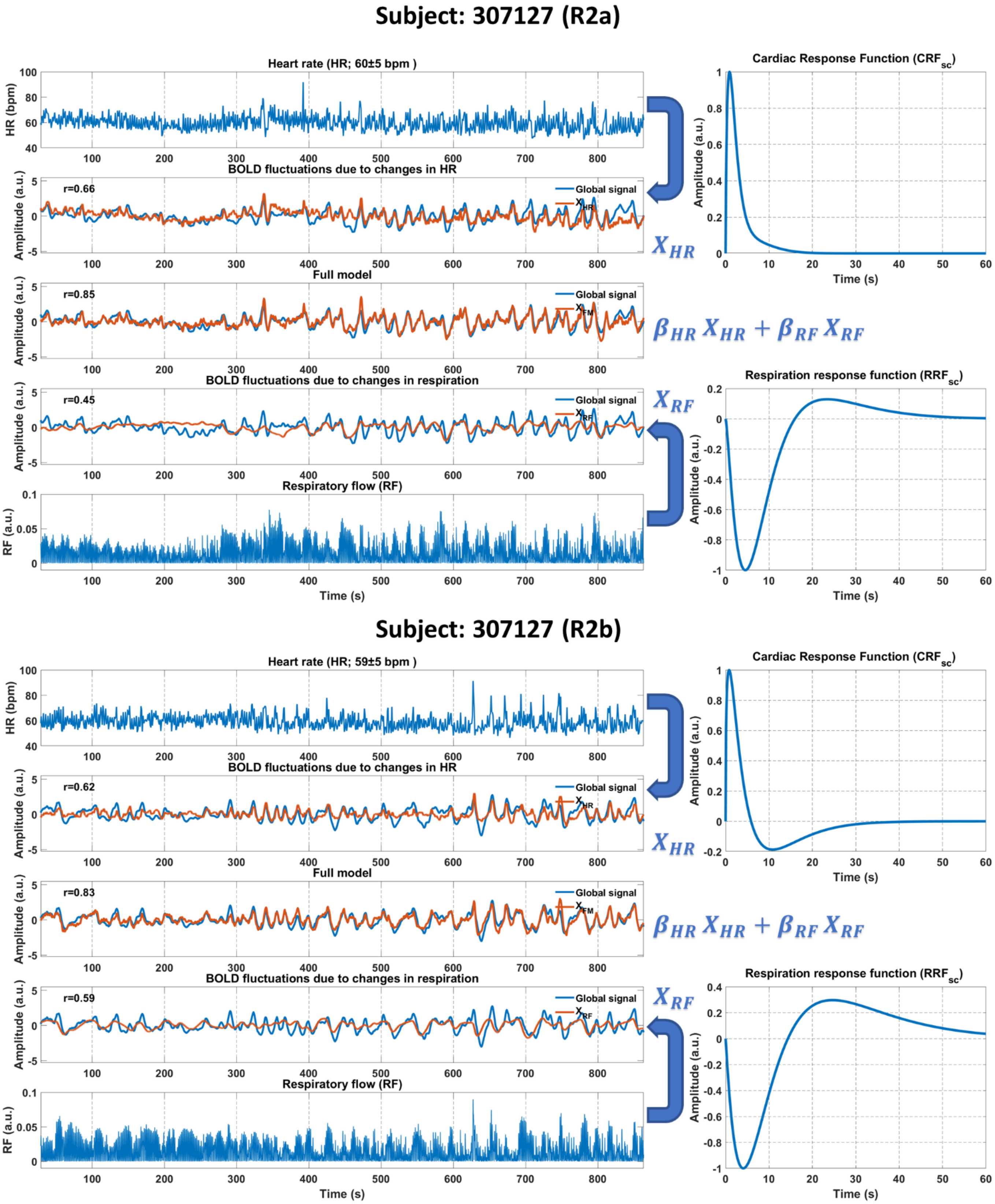


**Supplementary Fig. 7.** Demonstration of the $PRF_{sc}$ model for scans from the 2nd session of subject 307127

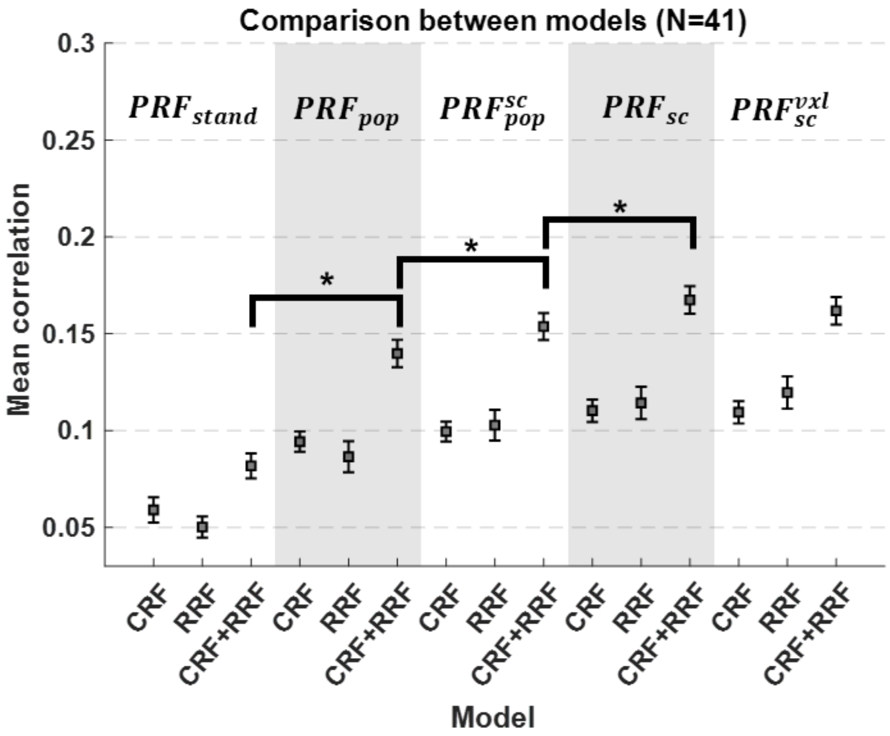


**Supplementary Fig. 8. Correlation values between physiological model predictions and voxel-specific timeseries, averaged over all voxels within a fixed ROI.** The squares and error bars indicate the mean and standard error of the means of all subjects. The ROI consisted of voxels with the 5% highest correlation in the group-level map for the $PRF_{stand}$ model. As in the case of the analysis with variable ROIs (Fig. 9), the scan-specific model $PRF_{sc}$ yielded the best performance. $*p < 10^{-6}$.

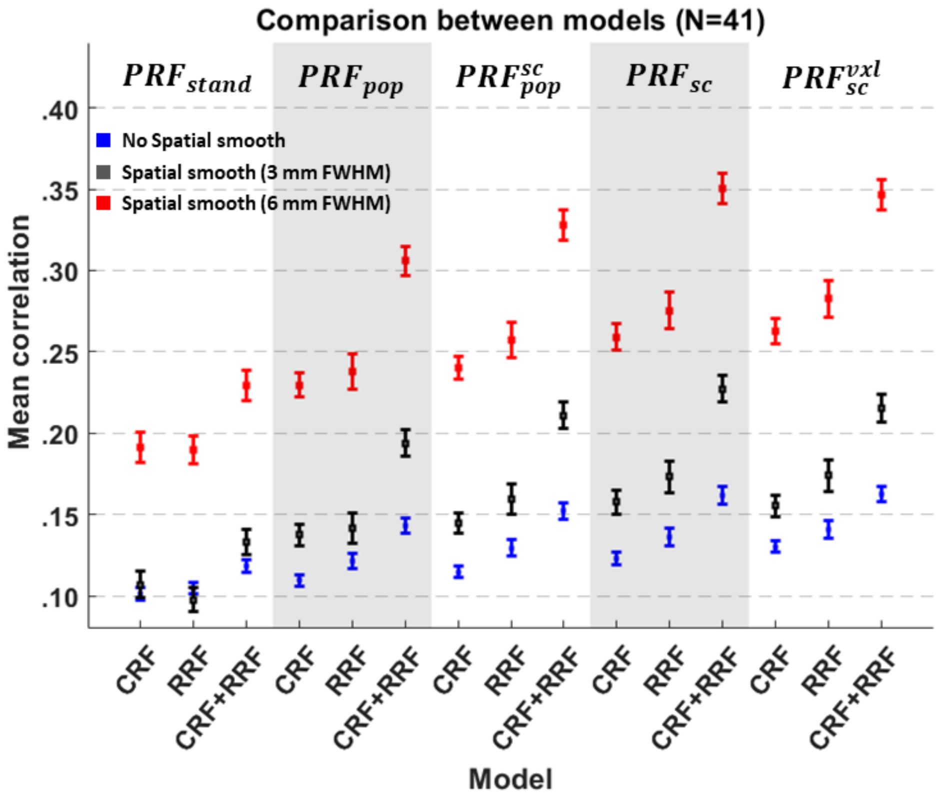


**Supplementary Fig. 9.** Correlation of the output of each model with the voxel timeseries in regions of interest (ROIs) for three different spatial smoothing filters. The squares and the error bars indicate the mean and standard error of the means of all subjects, and the color corresponds to different smoothing filters. Smoothing with 3 mm FWHM increased the mean correlation compared to not applying smoothing and, in turn, a wider filter of 6 mm FWHM increased even more the mean correlation. Note also that the effect of the spatial smoothing was the same in all models examined.

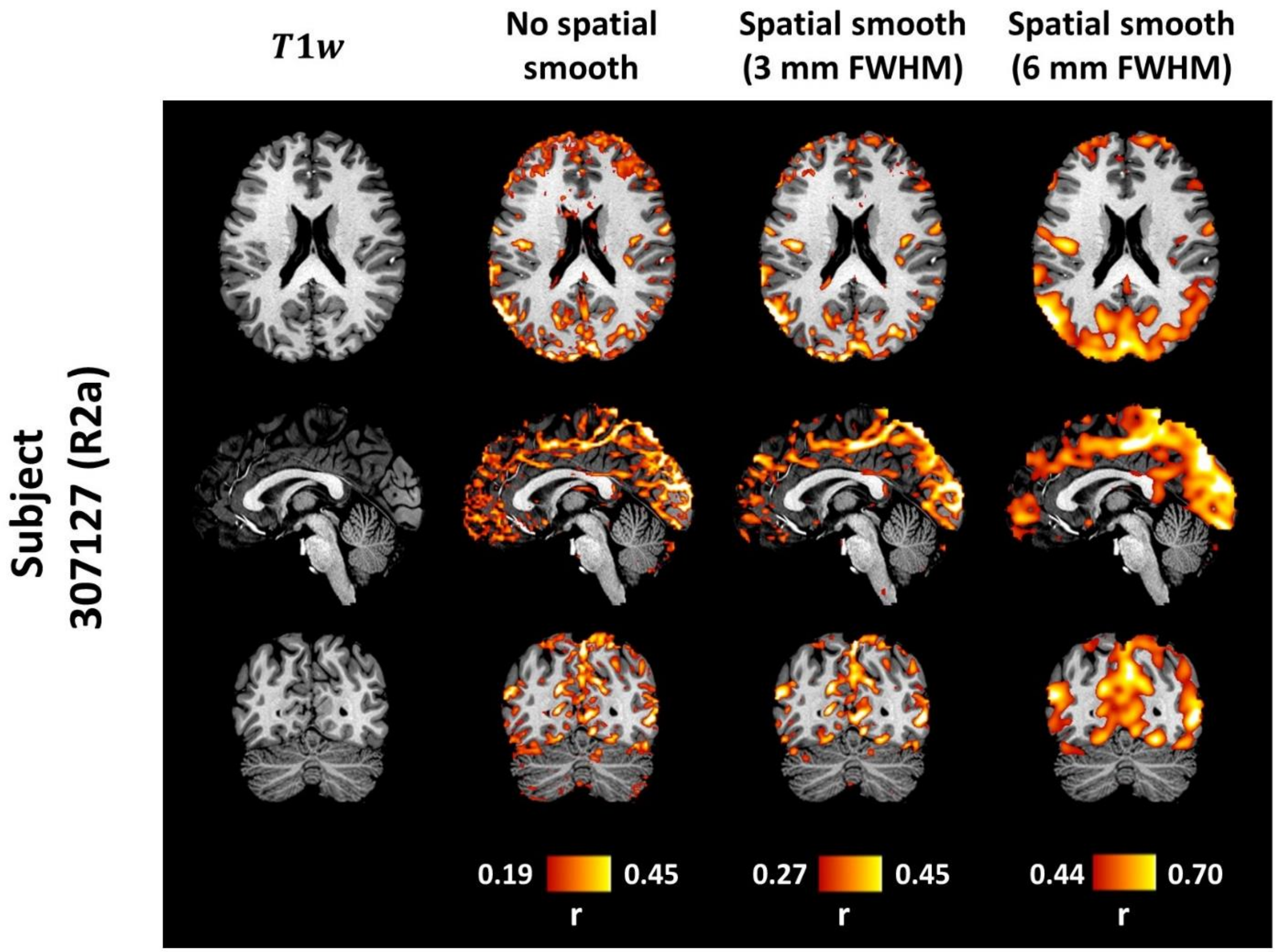


**Supplementary Fig. 10.** Maps of the correlation between the output of the $PRF_{sc}$ physiological model and fMRI timeseries for three different spatial smoothing filters. As in Supplementary Fig. 8, the wider was the spatial filter the higher were the correlation values. However, the 3mm FWHM filter retained much more spatial specificity compared to the 6 mm FWHM filter.

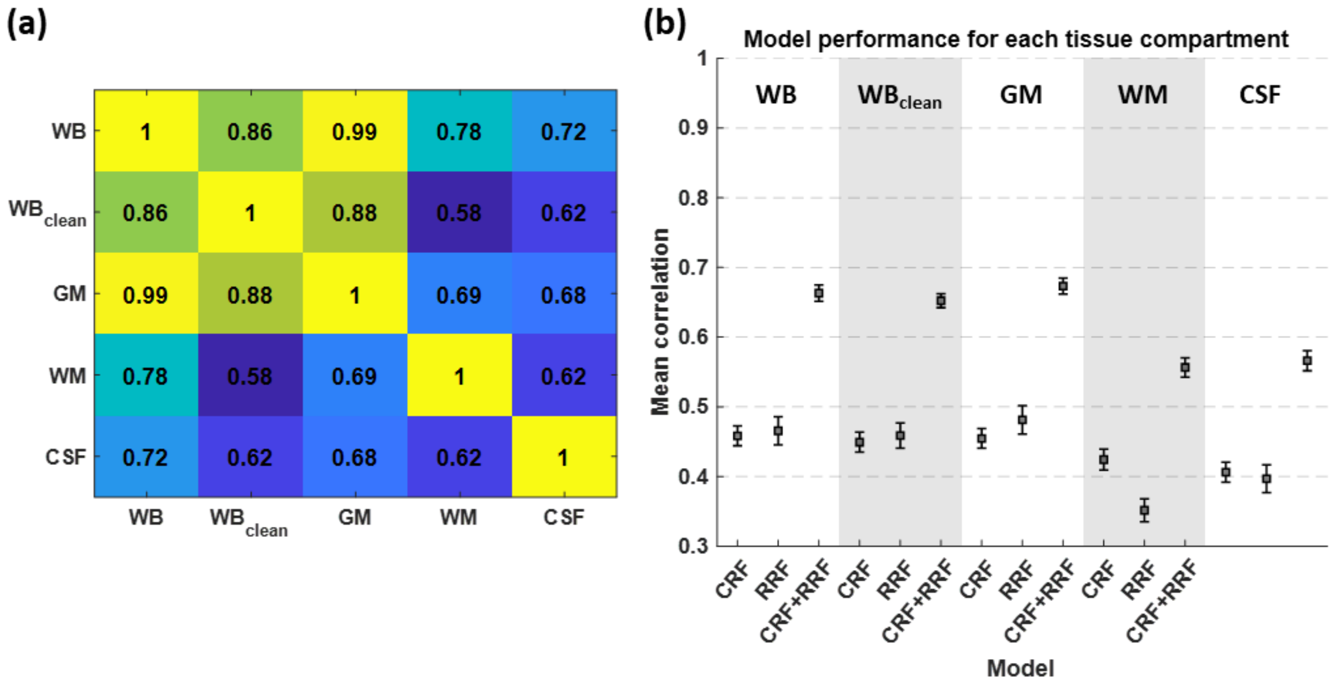


**Supplementary Fig. 11.** (a) Correlation matrix between mean time-series of tissue compartments averaged across scans and (b) model performance for each tissue compartment. Overall, we observe that the mean time-series from the three tissue type compartments, gray matter (GM), white matter (WM) and cerebrospinal fluid (CSF), are highly correlated (mean correlation ranges between 0.62-0.69). However, GM seems to contribute more to the whole brain (also termed as global signal (GS) in this work) as it is suggested by the mean correlation of 0.99 between these two timeseries. The effect of HR and RF is significantly more dominant in GM (and thus in GS) than in WM or CSF. Preprocessing before parameter estimation leads to a small decrease in the variance explained with the $PRF_{sc}$ model which suggests that there is small component on the GS explained by both $PRF_{sc}$ model and the nuisance regressors considered in the preprocessing (motion realignment parameters and high-frequency cardiac and respiratory regressors estimated with RETROICOR).

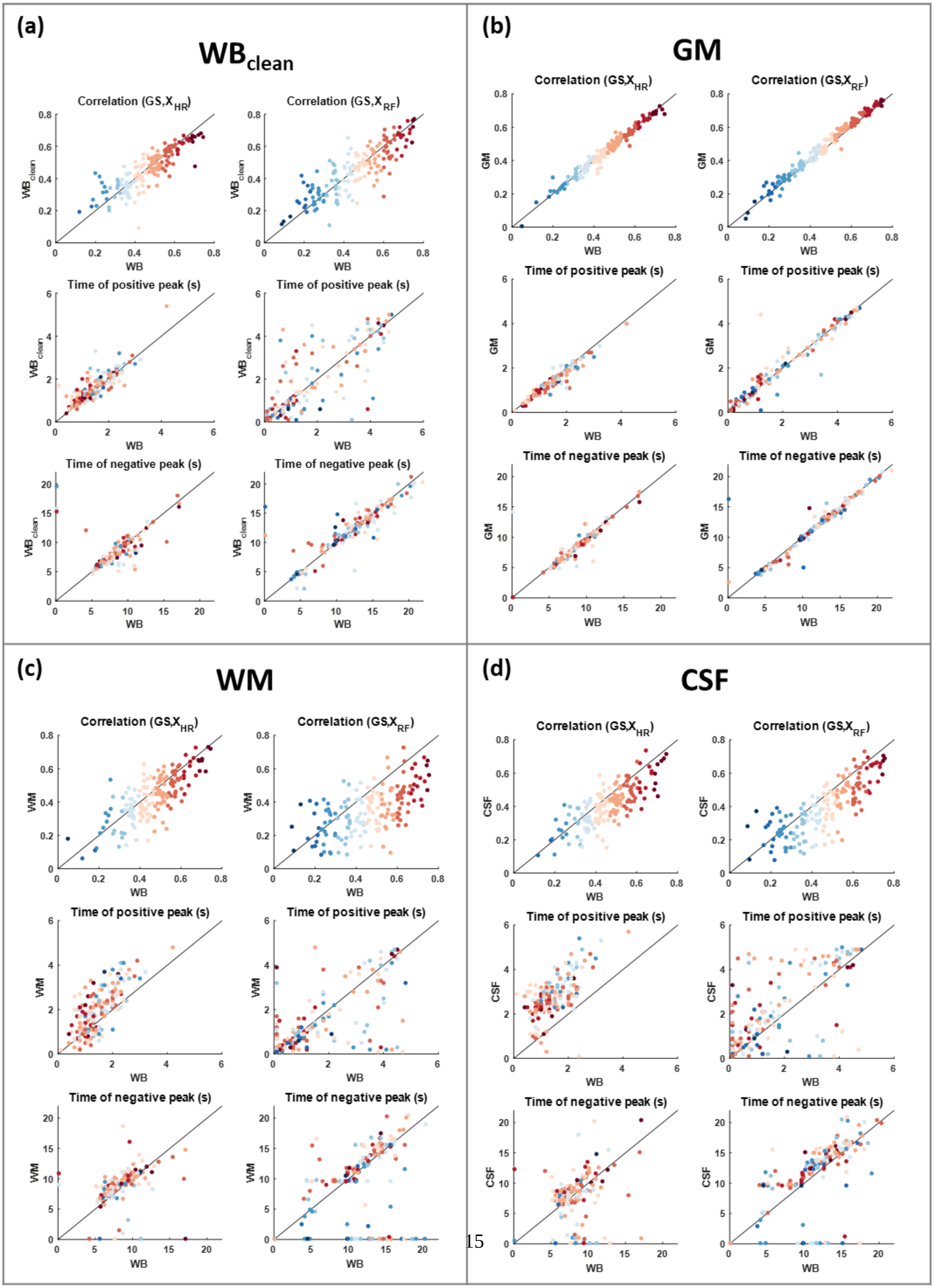

(a)
WB$_{clean}$
Correlation (GS,X$_{HR}$)
Correlation (GS,X$_{RF}$)
Time of positive peak (s)
Time of negative peak (s)
WB$_{clean}$
WB
(b)
GM
Correlation (GS,X$_{HR}$)
Correlation (GS,X$_{RF}$)
Time of positive peak (s)
Time of negative peak (s)
GM
WB
(c)
WM
Correlation (GS,X$_{HR}$)
Correlation (GS,X$_{RF}$)
Time of positive peak (s)
Time of negative peak (s)
WM
WB
(d)
CSF
Correlation (GS,X$_{HR}$)
Correlation (GS,X$_{RF}$)
Time of positive peak (s)
Time of negative peak (s)
CSF
WB

**Supplementary Fig. 12 (previous page).** Variability in the estimated $PRF_{sc}$ curves for different tissue compartment mean time-series. The $PRF_{sc}$ parameter estimation was performed using the five following dependent variables: the mean time-series from all voxels within the 1. whole brain (WB), 2. whole brain after regressing out nuisance regressors from each voxel ($WB_{clean}$), 3. gray matter (GM), 4. white matter (WM) and 5. cerebrospinal fluid (CSF). In all scatterplots, the x axis corresponds to the analysis with the WB mean time-series whereas the y axis in the four subplots corresponds to the analysis with the: (a) $WB_{clean}$, (b) GM, (c) WM and (d) CSF mean time-series. The variables represented in scatterplots correspond to the BOLD signal variance explained using HR and RF separately as well as to features regarding the shape of the $PRF_{sc}$ curves (e.g. time of positive peak). Overall, we observed that the explained variance and the $PRF_{sc}$ curve features were very similar for WB, $WB_{clean}$, GM and $GM_{clean}$. However, in the case of WM and CSF, the explained variance was significantly lower compared to WB, while the corresponding $PRF_{sc}$ curves exhibited overall slower dynamics (e.g. longer time of positive peak for $CRF_{sc}$).

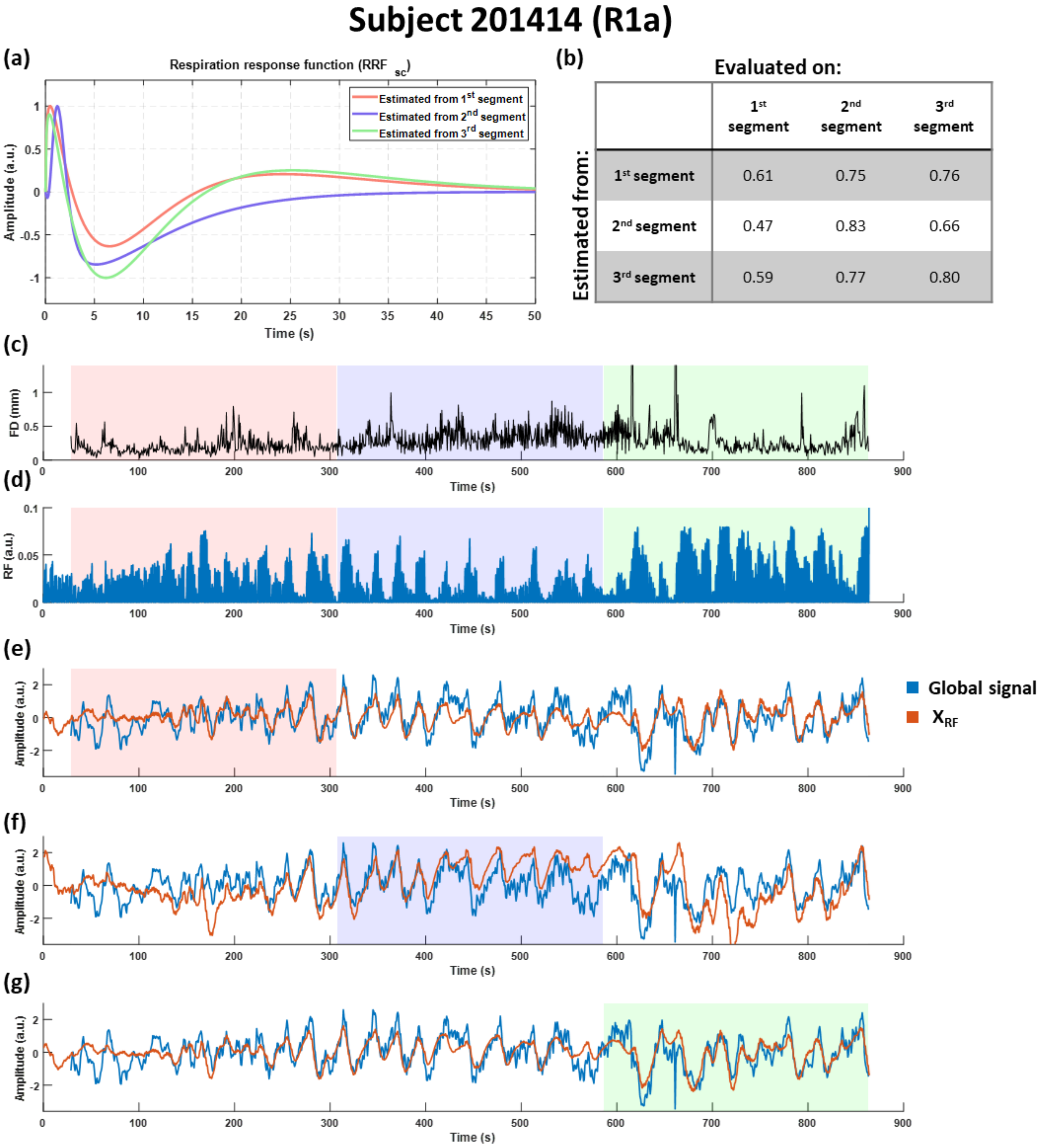


**Supplementary Fig. 13.** Variability in the estimated $RRF_{sc}$ curves for different time segments within a scan (Subject 201414 (R1a)). (a) Estimated $PRF_{sc}$ curves for the first, second and third 5-min segment within the scan. (b) Correlation between the GS and the model prediction for all nine possible combinations of training and validation segments. (c) Framewise displacement (FD) estimated from the motion realignment parameters as an index of head motion during the scan. (d) RF during the scan, where distinct patterns in breathing between the three time segments can be observed. (e-g) $RRF_{sc}$ model performance on the entire duration of the scan when the model parameters were estimated from the first, second and third segment, respectively. Overall, the $RRF_{sc}$ curves estimated from the three segments did not exhibit substantial differences despite the different breathing patterns observed between the segments.

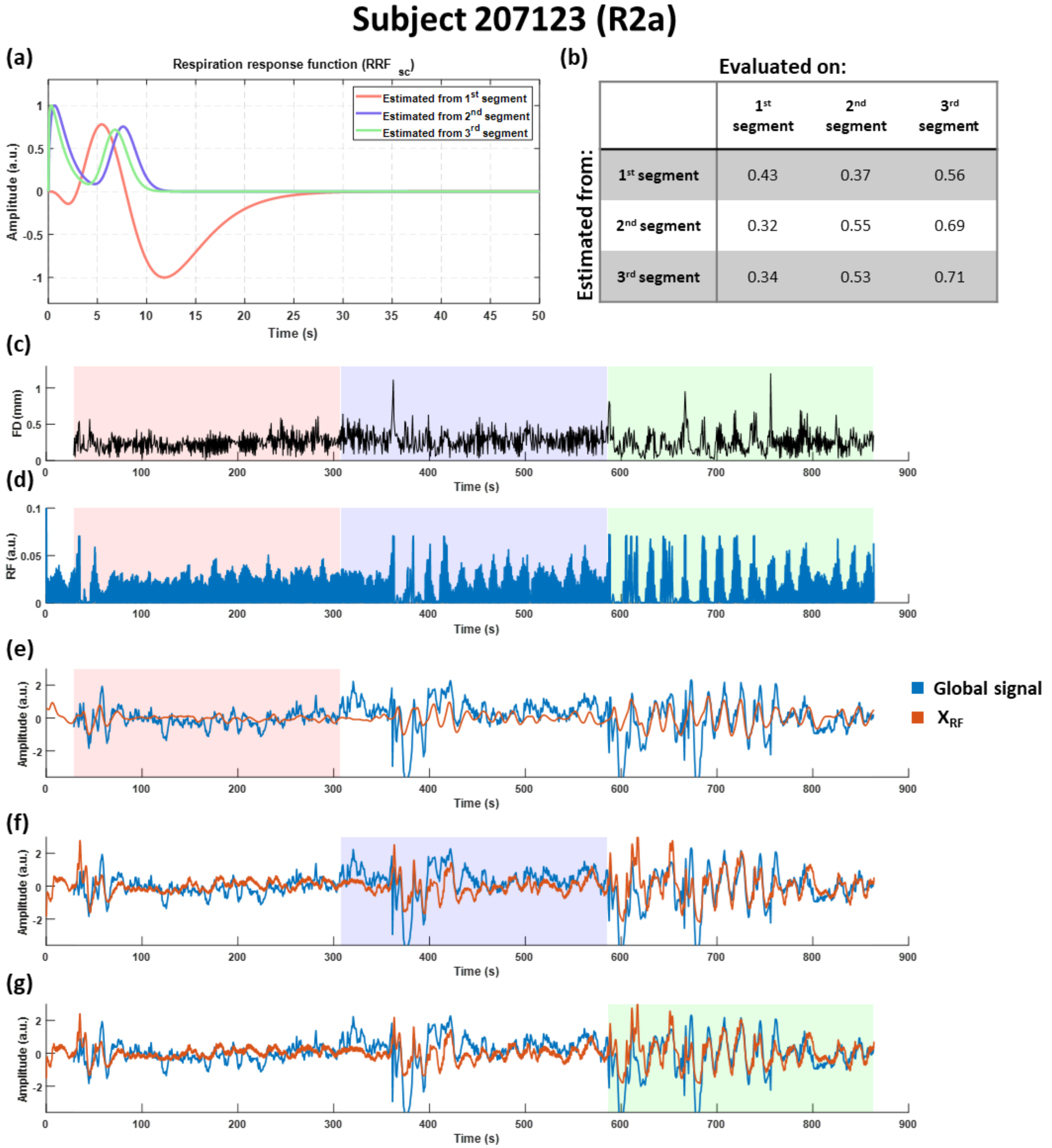


**Supplementary Fig. 14.** Variability in the estimated $RRF_{sc}$ curves for different time segments within a scan (Subject 207123 (R2a)). (a) Estimated $RRF_{sc}$ curves for the first, second and third 5-min segment within the scan. (b) Correlation between the GS and the model prediction for all nine combinations of training and validation segments. (c) Framewise displacement (FD) estimated from the motion realignment parameters as an index of head motion during the scan. (d) RF during the scan, where distinct patterns in breathing between the three time segments can be observed. (e-g) $RRF_{sc}$ model performance on the entire duration of the scan when the model parameters were estimated from the first, second and third segment, respectively. In this scan, the RF is fairly constant during the first segment indicating a stable breathing activity, while it is characterized by abrupt breathing patterns during the latter two segments. As a result, the $RRF_{sc}$ for the last two segments deviated significantly from the one estimated using the first segment. Importantly, as the RF fluctuates more within the third segment compared to the first segment, the variance explained in the third segment is higher than the variance explained in the first segment even when the $PRF_{sc}$ curve used in the model is the one estimated from the first segment.

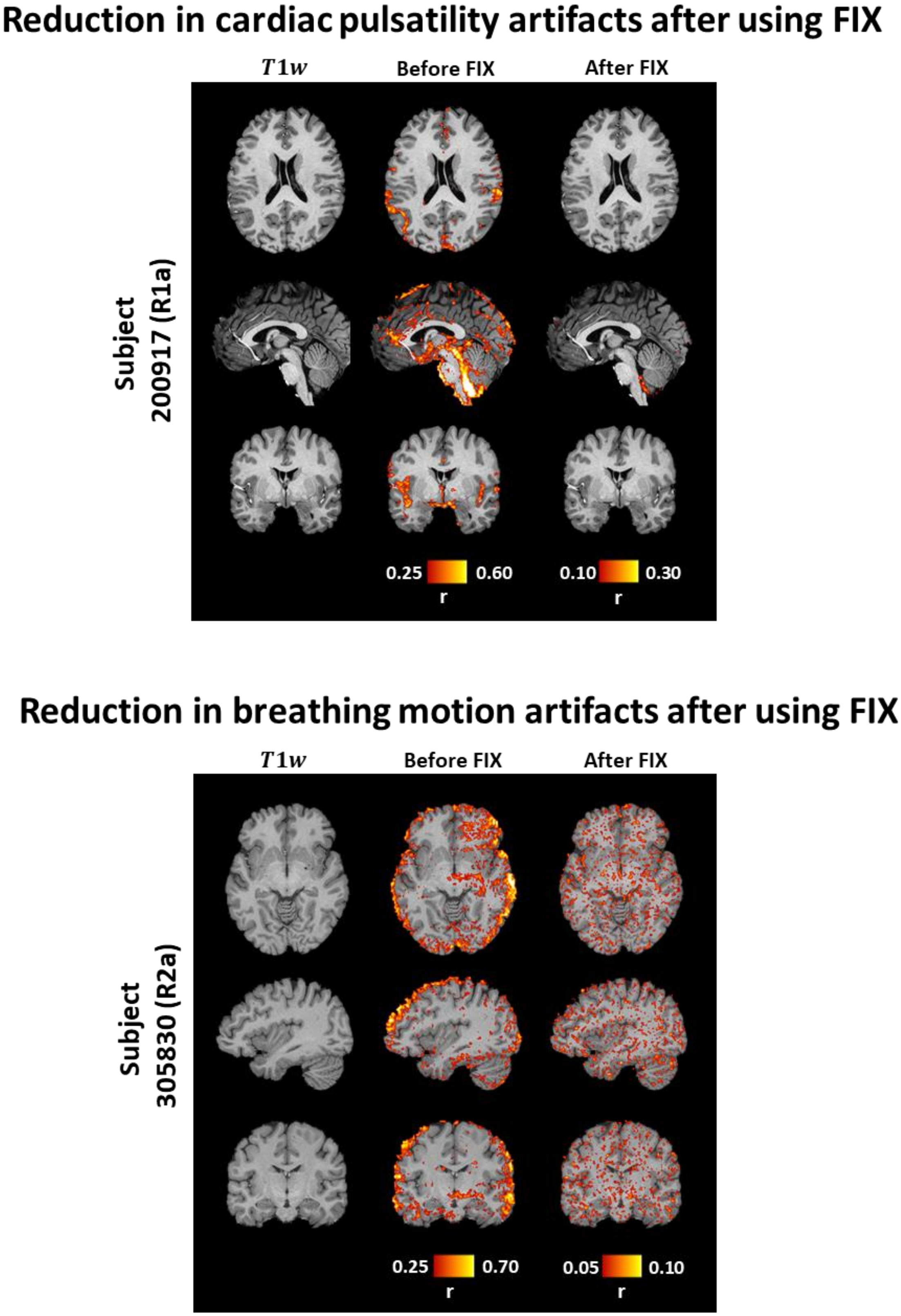


**Supplementary Fig. 15.** Correlation maps for representative scans related to cardiac pulsatility and breathing motion artifacts before and after using FIX. To extract the aforementioned maps, we used the cardiac- and breathing-related regressors in RETROICOR ($3^{rd}$ order) which are based on concurrent recordings of cardiac activity (photoplethysmogram) and breathing activity (respiratory belt). A significant reduction in cardiac pulsatility and breathing-motion artifacts was found after correcting the fMRI data using FIX.
